# MAIT cells protect in severe pneumococcal pneumonia by regulating neutrophil/macrophage antimicrobial activities

**DOI:** 10.1101/2025.02.07.637027

**Authors:** Emilie Barsac, Loïc Gonzalez, Guy Ilango, Antoine Hankard, Youenn Jouan, Roxane Lemoine, Vikrant Minhas, Marion Lenain, Matthieu Benard, Mégane Fernandez, Albanne Gaudio, Mustapha Si-Tahar, Antoine Guillon, Thierry Mallevaey, Charity Siu Gee Ganskow, Philipp Klahn, Jan-Willem Veening, Marion Salou, Olivier Lantz, Thomas Baranek, Christophe Paget

**Affiliations:** Inserm, Centre d’Etude des Pathologies Respiratoires (CEPR), UMR 1100, Tours, France; Université de Tours, Faculté de Médecine de Tours, France; Service de Médecine Intensive et Réanimation, CHRU de Tours, Tours, France; Service de Chirurgie Cardiaque et de Réanimation Chirurgicale Cardio-Vasculaire, CHRU de Tours, Tours, France; Unité Mixte de Service 61 « Analyses des Systèmes Biologiques », Département CSI, Université de Tours, CHRU de Tours, Inserm, Tours, France; Department of Fundamental Microbiology, Faculty of Biology and Medicine, University of Lausanne, Biophore Building, CH-1015 Lausanne, Switzerland; Department of Immunology, Temerty Faculty of Medicine, University of Toronto, Toronto, ON, Canada; Division of Organic and Medicinal Chemistry, Department of Chemistry and Molecular Biology, University of Gothenburg, Medicinaregatan 7B 413 90 Göteborg, Sweden; Center of Antibiotic Resistance Research in Gothenburg (CARe), Gulhedsgatan 10, 413 46 Göteborg, Sweden; Institut Curie, PSL University, Inserm U932, Immunity and Cancer, 75005 Paris, France

**Author notes:** Equal contribution (Thomas Baranek, Christophe Paget).

**Keywords:** MAIT, pneumococcus, neutrophil, bacterial clearance, lung

## Abstract

Mucosal-Associated Invariant T (MAIT) cells populate the lung tissue where they contribute to defense against respiratory infections. While MAIT cells have been implicated in host resistance to infections caused by Gram-negative bacteria, their contribution in immunity against Gram-positive bacteria-driven pneumonia is still enigmatic. Here, we demonstrate that both mouse and human MAIT cells are activated during severe infection caused by *Streptococcus pneumoniae,* the major cause of community-acquired bacterial pneumonia. Upon infection, lung MAIT cells undergo a transcriptional reprogramming associated with acquisition of potent antimicrobial properties. MAIT cell-deficient mice are more susceptible to pneumococcal pneumonia, including higher mortality, uncontrolled bacterial growth and dissemination, and impaired neutrophil and interstitial macrophage activity. Moreover, prophylactic stimulation of MAIT cells using cognate antigen protects from pneumococcus-induced lethal pneumonia. These findings demonstrate that MAIT cells are key cellular actors during Gram-positive bacterial infections.

## Introduction

*Streptococcus pneumoniae* (*S.p.*) is a Gram-positive bacterium that is often present in the upper respiratory tract. Although colonization by pneumococcus is usually associated with mild symptoms, it can become life-threatening^1^. Despite the availability of antibiotics and vaccines, *S.p.* remains one of the major cause of community-acquired pneumonia (CAP). In severe clinical presentations, pneumococcus-related pneumonia can lead to acute respiratory distress syndrome (ARDS) and sepsis requiring admission in intensive care unit (ICU) and mechanical ventilation^2^. In this context, recent estimates suggest that *S.p.*-associated diseases are responsible for at least half a million of deaths worldwide each year. Upon pneumococcus infection, the host immune response is rapidly mobilized to contain/eliminate the pathogen. During this early phase, alveolar macrophages (AM) and neutrophils are promptly recruited/activated to facilitate pneumococcus clearance through phagocytosis as well as the release of lytic enzymes, antimicrobial peptides and reactive oxygen species (ROS)^3^.

Several studies have highlighted the protective role of innate-like T (T_inn_) cells in experimental models of *S.p.*-induced pneumonia through their ability to regulate the activity of these phagocytes^4–7^. The underlying mechanisms rely on shared functions. For instance, γδT and invariant Natural Killer T (iNKT) cells produce large amounts of interleukin (IL)-17A that contribute to the early recruitment of neutrophils^4,5^. In addition, iNKT cells uniquely secrete granulocyte-macrophage colony-stimulating factor (GM-CSF), which is required for protection^7^. Moreover, increasing the pool of lung IL-17A-producing T_inn_ cells by means of IL-7 enables a better containment of *S.p.* in an experimental setting^6^. However, the relative contribution of Mucosal-Associated Invariant T (MAIT) cells, a third lineage of T_inn_ cells, in host defense to *S.p.* infection is unknown.

MHC-related protein 1 (MR1)-restricted MAIT cells mainly seed non-lymphoid tissues including barrier tissues, in which they exert versatile functions ranging from tissue physiology to immunosurveillance and tolerance^8–13^. As such, they orchestrate immune responses in a variety of contexts, including infection, cancer and inflammation. MAIT cells are characterized by the expression of a semi-invariant TCR composed of an invariant TCRα chain (TRAV1-2/TRAJ33 in humans and Trav1/Traj33 in mice) associated with TCRβ chains of limited diversity^14,15^. These highly conserved TCRs are specific to antigens (Ags) invisible to mainstream polymorphic MHC-restricted T cells. MAIT TCR recognizes metabolites generated from a non-enzymatic condensation of a microbial riboflavin pathway derivative (*e.g.* 5-A-RU) and small by-products of the glycolysis (glyoxal/methylglyoxal) to form pyrimidine antigens such as 5-OP-RU^16^. Thus, MAIT cells have emerged as critical components of the immune response to pathogens. Several studies have revealed the role of MAIT cells in immunity to respiratory bacterial infections induced by Gram-negative bacteria *Francisella tularensis*^17,18^*, Legionella longbeachae*^19^ and *Klebsiella pneumoniae*^20^. However, whether MAIT cells also contribute to the host response against Gram-positive bacteria such as *S.p.* remains unclear.

Interestingly, the riboflavin operon is highly conserved across *S.p.* serotypes, and several strains have been shown to activate MAIT cells *in vitro*^21,22^. Additionally, lung MAIT cells are rapidly activated *in vivo* upon *S. pneumoniae* infection and produce substantial levels of IL-17A^6,7^. However, using *Mr1*^-/-^ mice on a C57BL/6JRj (B6) background, MAIT cells were shown to be dispensable for pneumococcus clearance possibly due to redundant functions with γδT cells and iNKT cells^7^. Interestingly, local immunization with a MAIT ligand-producing bacterium (*e.g. Salmonella*) induces the expansion of two functional subsets of Ag-experienced MAIT cells expanded, characterized by expression of CD127 and Klrg1 and displaying a stable transcriptional and metabolic program associated with T_H_17- and T_H_1-like profile, respectively. Adoptive transfer of CD127^+^ Ag-experienced MAIT cells into B6 mice reduced the bacterial burden following *S. pneumoniae* infection^23^. Several clinical studies have also investigated MAIT cells in children and adults with bacterial CAP, including mild and severe forms^24–27^. These studies point towards a drastic decrease in circulating MAIT cells associated with altered activation status and cytokine profile. Moreover, this is paralleled with an increase proportion of activated MAIT cells in the airways^25,28^. Thus, these studies collectively suggest that MAIT cells may play relevant, yet undefined, functions in *S.p.* infection, and therefore may represent appealing targets for innovative anti-pneumococcal therapies.

Here, we show that ICU patients with severe forms of pneumococcal infections displayed a strong and uniform decrease in the proportion of circulating MAIT cells on admission. Importantly, activation levels of MAIT cells correlated with key clinical parameters of disease severity such as levels of hypoxemia. In response to experimental *S.p.*-induced pneumonia, lung MAIT cells were rapidly activated and acquired an antimicrobial transcriptional signature. Activation of MAIT cells was dependent on both cytokines and cognate signals. Moreover, MAIT cell-enriched mice exhibited higher protection to infection and better bacterial containment. This protective effect was associated with increased activity of key immune antibacterial players, including neutrophils and interstitial macrophages. Finally, prophylactic targeting of MAIT cells through instillation of a single dose of 5-OP-RU was sufficient to protect mice against lethal *S.p.*-induced pneumonia, a phenomenon associated with lung neutrophilia. These data demonstrate that MAIT cells are key cellular actors during Gram-positive bacterial infections.

## Results

### Activation status of circulating MAIT cells correlates with clinical features in patients with severe pneumococcal pneumonia

Human MAIT cells respond *in vitro* to various bacteria responsible for CAP^21,22,29^. Here, we analysed the frequency and phenotype of MAIT cells from peripheral blood mononuclear cells (PBMC) of nineteen adult patients upon ICU admission with microbiologically diagnosed bacterial CAP, and no other referenced pathogens (**Table 1**). As expected^30^, *Streptococcus pneumoniae* (*S.p.,* pneumococcus) was the most frequently identified pathogens in our patients (**Figure S1A**). The frequency of circulating MAIT cells within leukocytes from patients with severe bacterial pneumonia was decreased (**Figure 1A**), in agreement with previous studies^26,28^, and a greater proportion of their MAIT cells expressed the activation marker CD69 (**Figure 1B**) and the activation/exhaustion marker PD-1 (**Figure 1C**) with high variability, compared to healthy donors. A more marked decrease in MAIT frequency tended to be observed in patients with non-pneumococcal bacterial CAP (others) (**Figure S1B**). However, we could not observe differences in activation levels between groups (**Figure S1B**).

**Figure 1:**
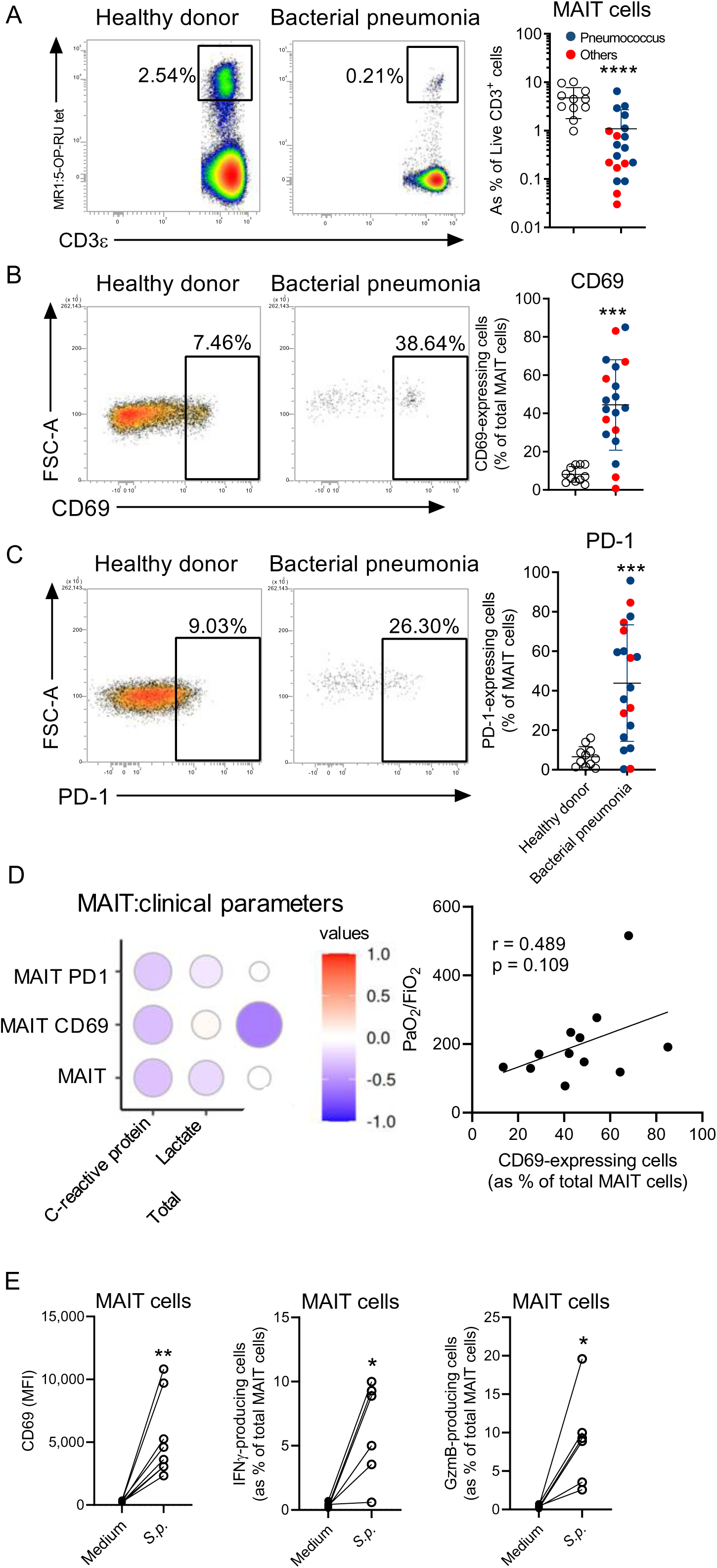
Phenotype of circulating MAIT cells and clinical features in ICU patients with severe bacterial pneumonia. (**A**), Representative dot plots of circulating MAIT cells (left panel) and MAIT cell frequency (right panel) from PBMC of ICU patients with severe bacterial pneumonia (n = 19) or healthy donors (n = 11) as controls. Mann-Whitney test. (**B-C**), Representative dot plots (left panels) and relative proportion (right panels) of CD69- (**B**) and PD-1-expressing (**C**) MAIT cells. Individual values and means ± SD are shown. Mann-Whitney test. (**D**), Spearman’s rank correlation of cell frequency, CD69 and PD-1 expression of MAIT cells with clinical features (left panel) and of PaO_2_/FiO_2_ ratio with CD69-expressing MAIT cells (right panel). SOFA : Sequential organ failure assessment (**E**) Activation of MAIT cells from PBMC of healthy donors cultured with fixed-*S. pneumoniae* serotype 1 (MOI 50). Wilcoxon test. *, P < 0.05; **, P < 0.01; ***, P < 0.001; ****, P < 0.0001.

To avoid pathogen-specific biases, we decided to focus on pneumococcal CAP (n = 12) for subsequent analyses. Since blood MAIT cell phenotypes have been shown to be correlated with SARS-CoV-2 infection outcome^31–33^, we assessed whether this was also true during severe pneumococcal CAP. The total sequential organ failure assessment (SOFA) score of the patients was negatively associated with high levels of CD69 on MAIT cells (**Figure 1D**). Interestingly, this was particularly true for the respiratory component of the SOFA since CD69 expression levels on MAIT cells positively correlated with the PaO_2_/FiO_2_ ratio (**Figure 1D**). This indicates lower hypoxemia in patients displaying highly activated MAIT cells, and therefore an association between MAIT cell phenotype and the respiratory functions. Finally, we tested whether MAIT cells from healthy donors could respond to a pneumococcal clinical isolate (E1586, serotype 1)^34^ used further for *in vivo* experimental settings in our study. Interestingly, MAIT cells from various donors were activated based on CD69 expression as well as production of IFN-γ and granzyme B (**Figure 1E**). Altogether, these data indicate that circulating MAIT cells from patients with severe pneumococcal CAP present activated phenotypes that can be associated with relevant clinical parameters.

### High content in MAIT cells confers protection against severe pneumococcal infection

Because our clinical data might suggest a role for MAIT cells in pneumococcal CAP, we decided to investigate MAIT cells in a model of severe pneumococcal infection^4,34^. The scarcity of MAIT cells in classical mouse laboratory strains constitutes a major impediment to preclinical investigation^35^. To circumvent this, we took advantage of congenic B6-MAIT^CAST^ mice, which have increased MAIT cells compared to B6 mice^35^ (**Figure 2A**). First, relative proportion of lung MAIT cells upon pneumococcal infection was not significantly modulated during the course of infection (**Figure 2A**). The frequency of CD69^+^ lung MAIT cells significantly increased from 12 hours post-infection (hpi) in B6-MAIT^CAST^ mice, and from 24 hpi in B6 mice (**Figure 2B**). Of note, PD-1 expression by lung MAIT cells remained unchanged over time (**Figure 2B**).

**Figure 2:**
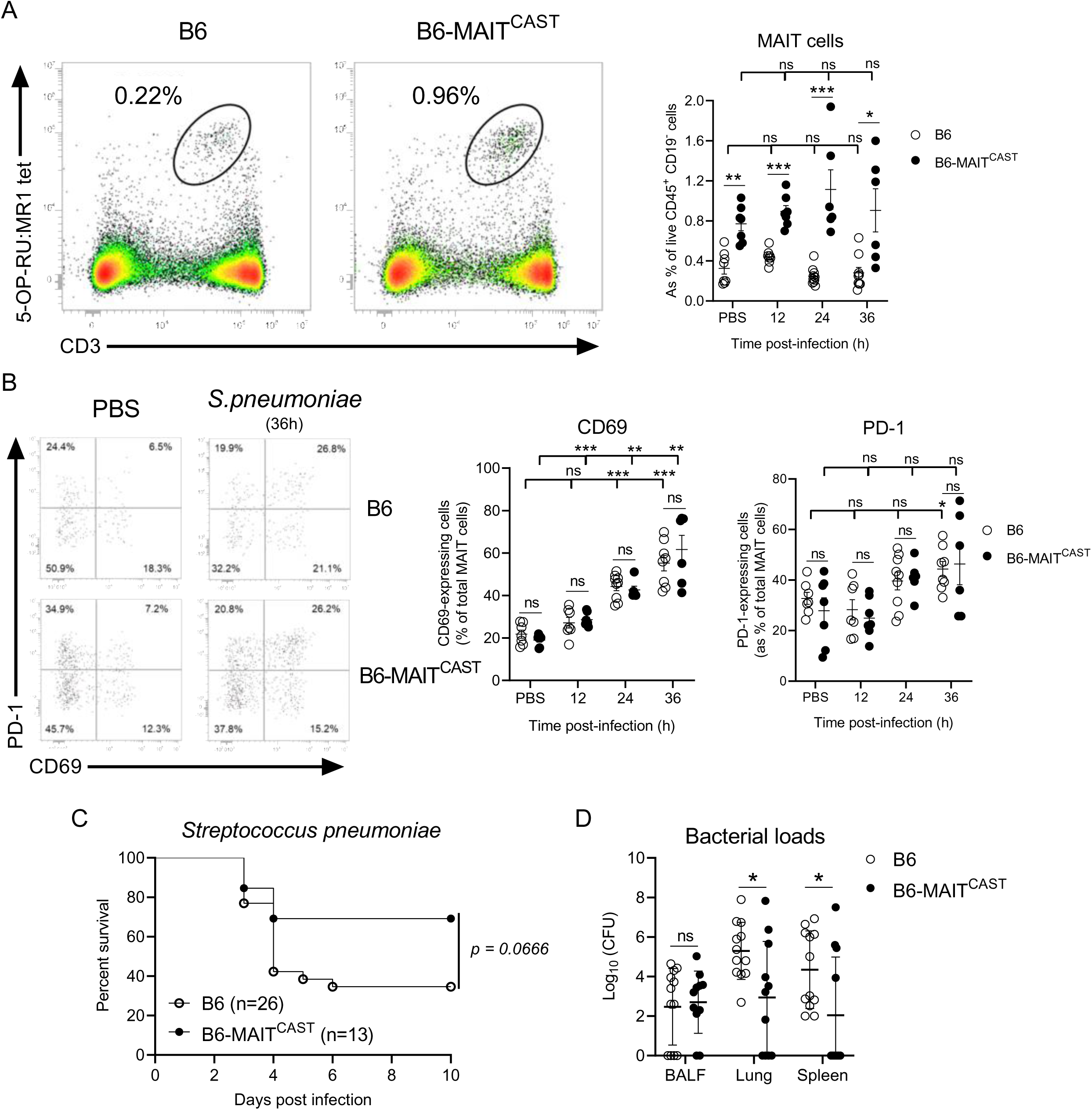
MAIT cells contribute to the anti-pneumococcal defense. (**A-D**) B6 and B6-MAIT^CAST^ mice were i.n. infected with *S. pneumoniae* serotype 1 with 2 x 10^6^ pfu (**A-B**) or 5 x 10^5^ pfu (**C-D**). (**A-B**) Mice were euthanized at the indicated time point. (**A**), Frequency of lung MAIT cells in B6 and B6-MAIT^CAST^ mice during the course of infection. Representative dot plots (left panel), and individual values and means ± SD (right panel) are shown. Data are pooled from three independent experiments. Mann-Whitney test. (**B**), MAIT cell activation based on CD69 and PD-1 expressions was determined by flow cytometry. Representative dot plots (left panel), and individual values and means ± SD (right panel) are shown. Data are representative of two independent experiments. Mann-Whitney test. (**C**), Survival of mice was monitored daily (n = 13-26/group). Data are pooled from two independent experiments. Mantel-Cox test. (**D**), Bacterial burden was evaluated in broncho-alveolar lavage fluids (BALF), lung parenchyma and spleen at 60 hpi. Data are pooled from two independent experiments. Individual values and mean ± SD are shown. Mann-Whitney test. ns, not significant; *, P < 0.05; **, P < 0.01 ; ***, P < 0.001; ****, P < 0.0001.

Interestingly, while ∼ 60% of B6 mice succumbed within 4 days of infection, this was the case for only 30% of B6-MAIT^CAST^ mice (**Figure 2C**), although the difference did not reach significance. These data suggested that MAIT cell numbers in mice may correlate with host resistance to pneumococcus. Accordingly, the higher resistance of B6-MAIT^CAST^ mice was associated with a better control of local bacterial growth and systemic dissemination (**Figure 2D**). These data indicate that MAIT cells contribute to the resistance to pneumococcal infection and to the bacterial clearance.

### *S. pneumoniae* infection leads to a transcriptional reprogramming of lung MAIT cells associated with potent antibacterial activities

To explore the transcriptional landscape of MAIT cells during the course of pneumococcal infection, sorted MR1:5-OP-RU-specific lung T cells from naive or *S.p.*-infected (12 and 36 hpi) B6-MAIT^CAST^ mice were subjected to droplet-based single-cell RNA-sequencing (**Figure S2A**). Upon quality controls and filtering steps, 13,116 cells (PBS: 3,157; 12hpi: 7,441 and 36hpi: 2,518) were considered for downstream analyses. The datasets were merged and unsupervised cell clustering was performed^36^, enabling the identification of eight clusters (**Figure 3A**). Based on the cardinal transcription factors *Zbtb16*, *Tbx21* and *Rorc*, cluster 6 was identified as MAIT1 while clusters 1-5 and 7 comprised MAIT17 cells (**Figure 3B**). Cluster 8 cells lacked expression of these three transcription factors (**Figure 3B**) but expressed genes associated with a circulating phenotype (*Klf2, Ccr7, Sell*) (**Figure S2B and Table S1**) that suggests a cluster of MR1:5-OP-RU-specific naive-like T cells^37^ or contaminant conventional T cells and, therefore was not considered for further analyses. Confirming our assignment, differentially expressed genes (DEG) in cluster 6 comprised *Ccl5*, *Nkg7*, *Gramd3* and *Gimap4* (**Figure S2B and Table S1**) that are highly related to the MAIT1 program^38^. In line with *Rorc* expression, clusters 1-5 and 7 shared expression of genes associated with MAIT17 biology such as *Lgals3*, *Icos*, *Tmem176a* and *Tmem176b* (**Figure S2B and Table S1**). Within MAIT17 clusters, cluster 7 exhibited a unique signature featuring interferon-stimulated genes (ISG) such as *Stat1*, *Isg15*, *Bst2* and *Cxcl10* (**Figure S2B and Table S1**) and, therefore, was referred to as MAIT17-ISG.

**Figure 3:**
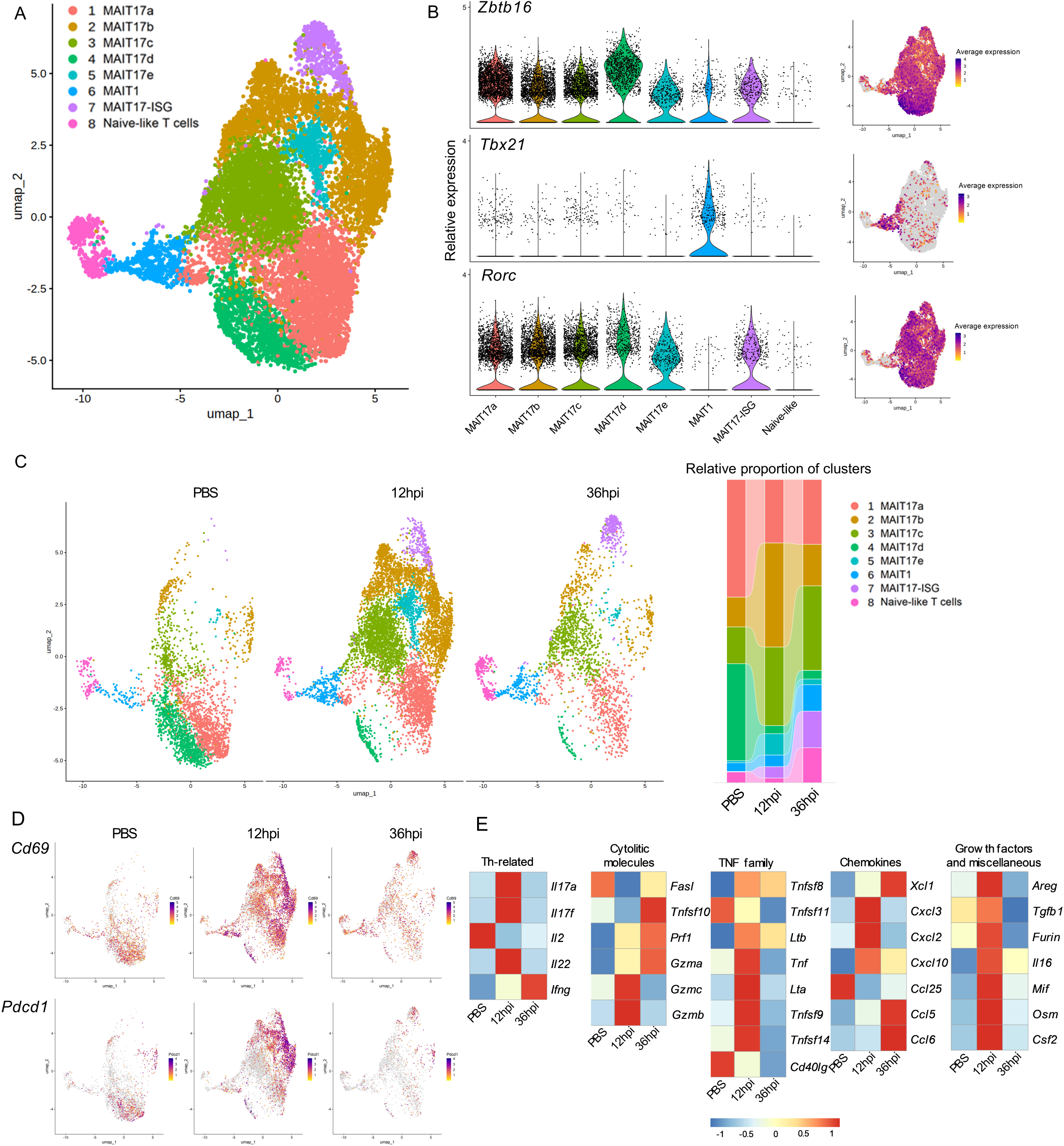
scRNA-seq profiling of lung MAIT cells from naive and *S.p*.-infected mice. B6-MAIT^CAST^ mice were infected or not with 5 x 10^5^ pfu of *S. pneumoniae* serotype 1 and euthanized at indicated time points. Lungs were collected and processed prior MAIT cell sorting (CD45^+^ CD19^-^ CD3^+^ MR1:5-OP-RU tetramer^+^). (**A**), Identification of cell clusters using the graph-based Louvain algorithm (resolution = 0.2) on Uniform Manifold Approximation and Projections (UMAP). Each dot represents one cell. (**B**), Expression of *Zbtb16*, *Tbx21* and *Rorc* per cluster is shown in the left panel and projection on aggregated UMAP is shown in the right panel. (**C**), Identity of clusters per condition on aggregated UMAP is depicted in the left panel. Relative proportions of each cluster per condition is shown in the right panel. (**D**), Projection of *Cd69* (upper panel) and *Pdcd1* (lower panel) on aggregated UMAP per condition is shown. (**E**), Heat maps of T cell-derived mediators expressed in MAIT transcriptomes per condition (pseudo-bulk) are shown. Mediators have been categorized per family.

Partition of clusters according to hpi indicated that the proportion of MAIT1 was not modulated during the early course of infection (12 hpi) but slightly increased at 36 hpi (**Figure 3C**). Within MAIT17, the relative proportion of clusters was highly modulated upon infection with some clusters repressed (MAIT17a and MAIT17d) and other being increased (MAIT17b and MAIT17c). Interestingly, MAIT17e and MAIT17-ISG were induced by *S. pneumoniae* infection (**Figure 3C**). Cluster proximity indicated that MAIT17 clusters could be categorized in three groups (**Figure S2C**) related to the infection status: Group 1 comprised MAIT17a and MAIT17d (mainly from control mice) and group 2 comprised MAIT17b, MAIT17e and MAIT17-ISG (mainly MAIT cells from *S.p.*-infected mice). Notably, MAIT17c (group 3) presented the highest divergence with the other MAIT17 clusters (**Figure S2C**). Thus, these data demonstrate an unappreciated transcriptional heterogeneity in lung MAIT cells especially within MAIT17, which is dynamically modulated upon pneumococcal infection.

Based on gene expression, we observed that pneumococcal infection led to lung MAIT cell activation (*Cd69*, *Pdcd1*) (**Figure 3D and S2D**). Consistent with the FACS data, highest expression was mainly found in pneumococcus-induced clusters (**Figure S3D**). We defined a panel of ∼100 genes encoding for acknowledged T cell-derived mediators involved in anti-infective response (**Table S2**). Notably, pseudo-bulk analysis revealed pneumococcus-driven regulation of 33 DEG linked to T_H_-related response, cytotoxicity, TNF family, chemokines, growth factors, and miscellaneous factors (**Figure 3E and Table S2**).

Altogether, these data indicate that pneumococcus infection remodels lung MAIT cell transcriptional program towards antibacterial and cytotoxic functions.

### Cytokines and TCR engagement are required for MAIT cell activation during *S. pneumoniae* infection

We next characterized the activation mechanisms of lung MAIT cells in response to pneumococcus infection. Given the evidence of the conservation of the riboflavin operon in most pneumococcal serotypes^21^, we first evaluated the contribution of TCR signals using *Nr4a1*-GFP B6-MAIT^CAST^ mice^39^. Interestingly, while activation of MAIT cells could be observed as soon as 12 hpi (**Figure 2B**), increased expression of GFP was only detected at 36 hpi in MAIT cells from the lungs of *S.p.*-infected mice (**Figure 4A**). Of note, this late TCR triggering is associated with the increase of the bacterial load (**Figure S3A**), possibly leading to higher bioavailability of riboflavin metabolites at 36 hpi. To confirm the involvement in TCR signals, we generated a mutant strain with a selective deletion in the entire riboflavin operon (*S.p.* Δ*RibDEHA*), which is required to produce the MAIT Ag precursor 5-A-RU. As compared to control strains, we observed a partial but significant decrease in CD69 expression on MAIT cells from mice infected with *S.p.* Δ*RibDEHA* (**Figure 4B**). However, deletion of the riboflavin operon did not have any effect on the ability of lung MAIT cells to produce IL-17A (**Figure 4C**). Further, cultures of human PBMC from healthy donors in presence of *S.p.* increased expression of CD69 on MAIT cells, an effect that was partially abrogated using the *S.p.* Δ*RibDEHA* strain (**Figure 4D**). Notably, no clear differences in bacterial growth between parental and mutant strains were observed *in vitro* (**Figure S3B**).

**Figure 4:**
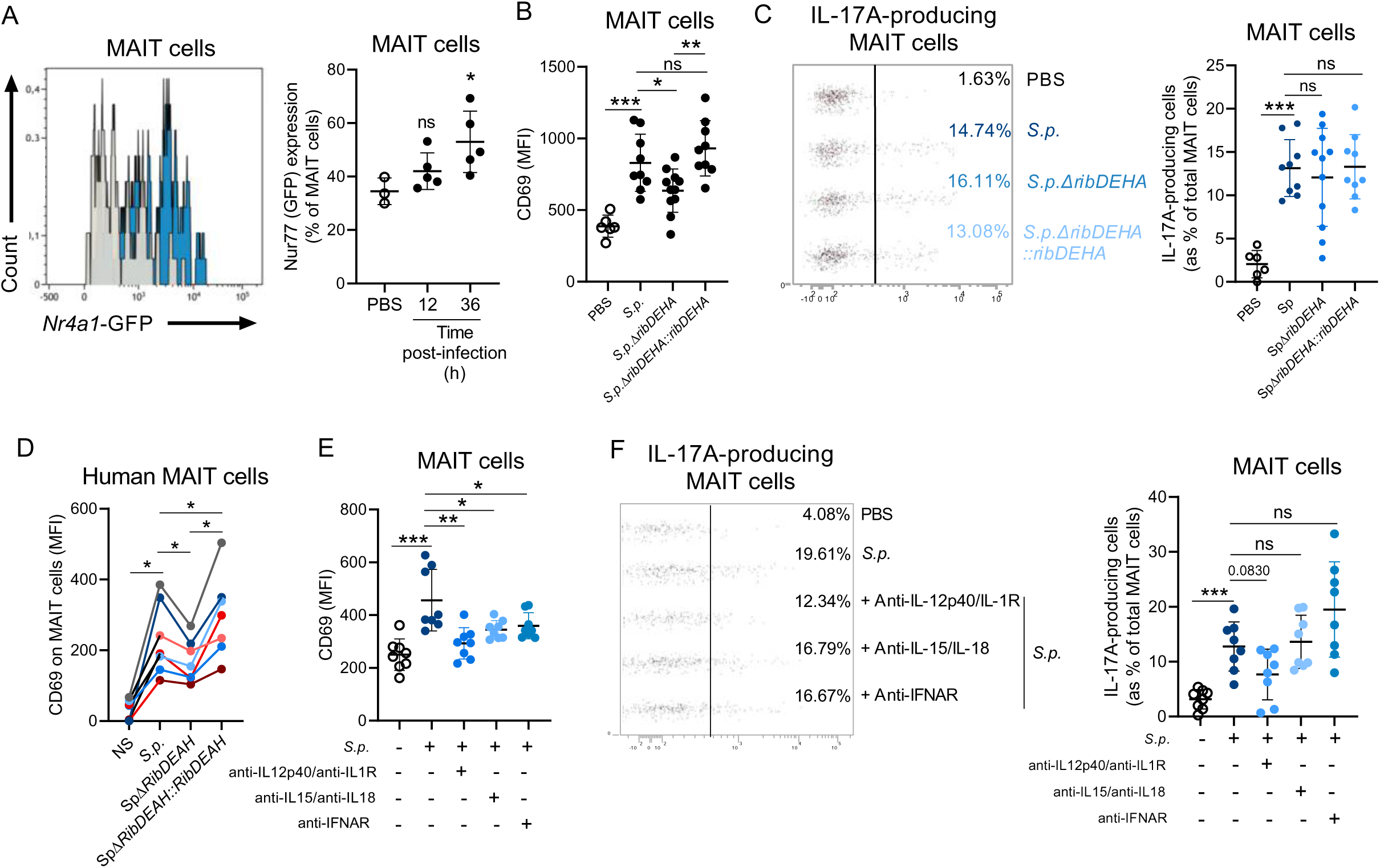
Activation mechanisms controlling MAIT cell activation during pneumococcal infection. (**A**) *Nr4a1*-GFP mice were i.n. infected with *S. pneumoniae* serotype 1 (2 x 10^6^ pfu). Lung MAIT cell activation was evaluated by analyzing the GFP expression under the control of *Nr4a1* (Nur77). Representative histograms (Grey: PBS; Blue: 36hpi) are shown in left panel and individual values and mean ± SD mean are depicted in the right panel. Data are representative of two independent experiments. Mann-Whitney test. (**B, C**) B6 mice were i.n. infected with 4 x 10^6^ pfu of *S. pneumoniae* D39V parental (*S.p.*) and mutant strains deficient for 5-A-RU production (*S.p.* Δ*RibDEHA*) or recomplemented (*S.p.* Δ*RibDEHA::RibDEHA*), and activation of lung MAIT cells was analyzed at 24 hpi by flow cytometry. (**B**), Individual values and mean ± SD of CD69 expression from two independent experiments are depicted. Mann-Whitney test. (**C**), Representative dot plots of the frequency of IL-17A-expressing MAIT cells are shown in the left panel. Individual values and mean ± SD from two independent experiments are shown in the right panel. Mann-Whitney test. (**D**), Fixed-pneumococcal strains (MOI 50) were cultured with PBMC from healthy donors and CD69 expression on MAIT cells was determined after 12h of culture. Individual values of CD69 expression from seven donors are shown. Wilcoxon test. (**E-F**), B6 mice were i.n. treated or not with combination of monoclonal Abs, and infected with *S. pneumoniae* serotype 1 (2 x 10^6^ pfu) the following day. Expression of CD69 (**E**) and intracellular IL-17A (**F**) by MAIT cells were evaluated at 24 hpi. (**E**), Individual values and mean ± SD of CD69 expression on MAIT cells from two independent experiments is shown. Mann-Whitney test. (**F**), Representative dot plots of the frequency of IL-17A-expressing MAIT cells are shown in the left panel. Individual values and mean ± SD from two independent experiments are shown in the right panel. Mann-Whitney test. ns, not significant; *, P < 0.05; **, P < 0.01; ***, P < 0.001.

Since disruption of the MR1:Ag/TCR interaction only partially affected MAIT cell activation, we hypothesized that other signals, such as cytokines, may participate in MAIT cell activation. Thus, the potential contribution of these cytokines in MAIT cell activation was biologically evaluated *in vivo* using neutralizing/blocking antibodies. Blockade of the IL-1β/IL-12p40, IL-18/IL-15 and type I IFN pathways led to a decrease in MAIT cell activation as judged by CD69 expression (**Figure 4E**). However, only IL-1β/IL-12/IL-23 receptor signaling neutralization tended to decrease IL-17A production albeit in a non-significant manner (**Figure 4F**). Thus, these data indicate that pneumococcus activates MAIT cells via both cytokine and TCR-mediated signals.

### MAIT cell protection against *S. pneumoniae* infection is associated with higher antibacterial functions of neutrophils and interstitial macrophages

As our data suggested that MAIT cell numbers correlated with host resistance to *S. pneumoniae* infection, we next compared the susceptibility of B6-MAIT^CAST^ and B6-MAIT^CAST^ *Mr1*^-/-^ mice, which lack MAIT cells. As expected, MAIT cell-deficient mice were more susceptible to infection as compared to B6-MAIT^CAST^ (**Figure 5A**). Of note, survival rate of B6-MAIT^CAST^ *Mr1*^-/-^ mice was comparable to that of B6 mice (**Figure 2C)**. Moreover, bacterial burden was significantly increased in the airways and lung parenchyma of B6-MAIT^CAST^ *Mr1*^-/-^ infected mice (**Figure 5B**). Further, we noticed an increased bacterial dissemination in MAIT cell-deficient mice as assessed by bacterial loads in the spleen (**Figure 5B**). Transcriptome of lung MAIT cells from *S.p.*-infected mice displayed a gene signature previously associated to acute lung infection^40^ at 12 hpi (**Figure 5C**) with highest expression in clusters (**Figure S4A**) belonging to group 2 (**Figure S2C**). Of note, transcriptomes of MAIT cells from infected mice also harbored a higher “tissue repair” signature^41^ at 12 hpi (**Figure S4B**). Since containment of *S.p.* is largely mediated by IL-17A-dependent recruitment of neutrophils in this model^5,6^, we assessed neutrophil infiltrate in the lungs. We observed a decrease in the relative proportion of lung - but not BALF - neutrophils from infected B6-MAIT^CAST^ *Mr1*^-/-^ mice compared to B6- MAIT^CAST^ mice (**Figure S4C**). However, no changes in absolute numbers were observed (**Figure 5D**). This absence of clear phenotype may be due to overlapping functions with γδT17 cells^7^ that are numerically more abundant than MAIT17 cells (**Figure S4D**). Analysis of other relevant effector cells in pneumococcus containment such as alveolar macrophages (AM) (CD11b^-/dim^ SiglecF^+^) and interstitial macrophages (CD11b^+^ SiglecF^-^ Ly6G^-^ F4/80^+^) did not indicate changes in absolute numbers either (**Figure S4E**).

**Figure 5:**
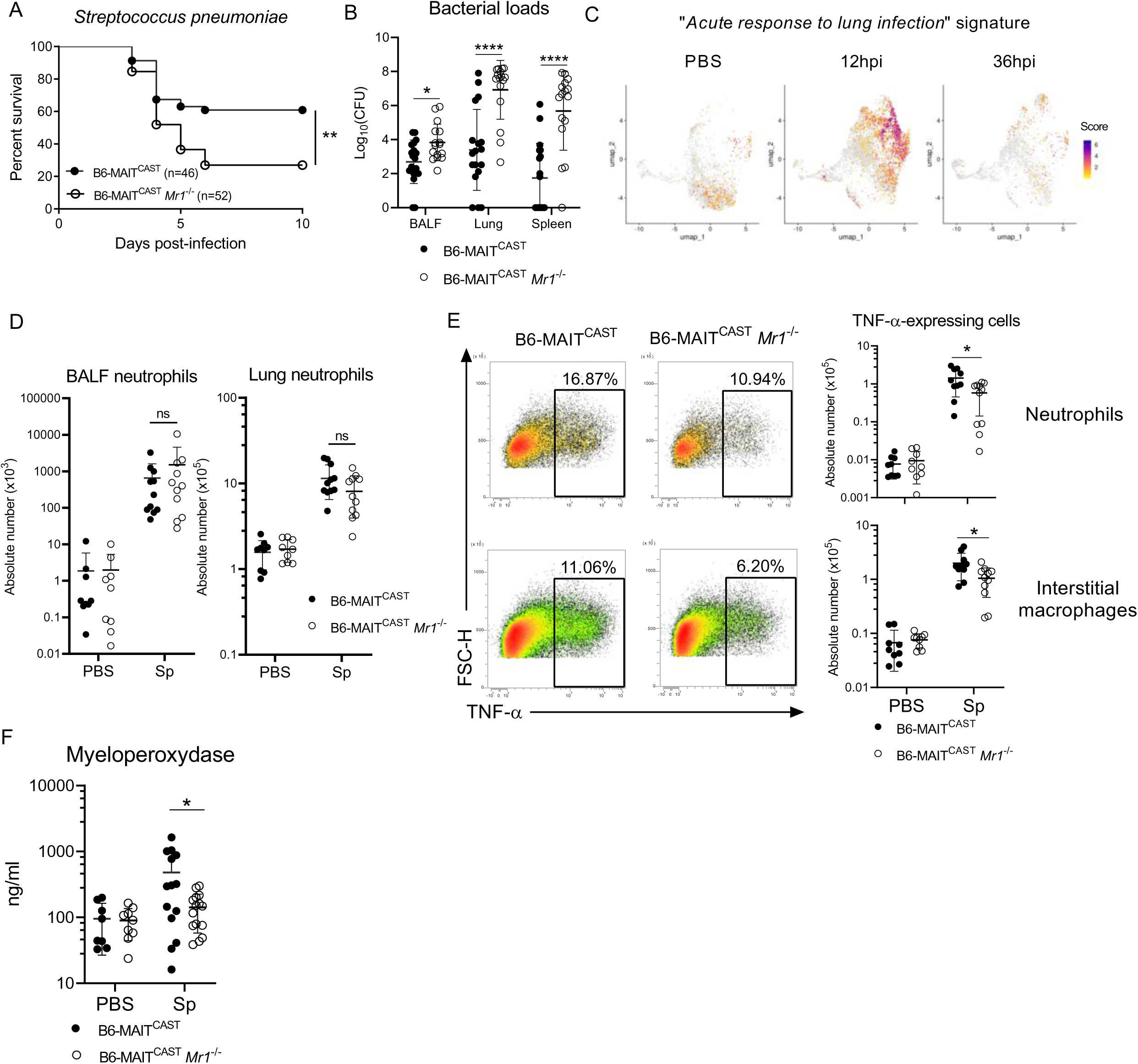
Influence of MAIT cells on phagocyte activity during pneumococcal infection. (**A-C**) B6-MAIT^CAST^ and B6-MAIT^CAST^ *Mr1*^-/-^ mice were i.n. infected with *S. pneumoniae* serotype 1 (5 x 10^5^ pfu). (**A**), Survival of mice was monitored daily for 10 days. Data are representative of seven independent experiments. Mantel-Cox test. (**B**), Individual values and mean ± SD of bacterial loads (60 hpi) in indicated compartments from four independent experiments are depicted. Mann-Whitney test. (**C**), Evaluation of the “*Acute response to lung infection*” signature on aggregated lung MAIT transcriptomes from naive and 12 hpi and 36 hpi *S. pneumoniae*-infected B6-MAIT^CAST^ mice. (**D-E**) B6-MAIT^CAST^ and B6-MAIT^CAST^ *Mr1*^- /-^ mice were i.n. infected with *S. pneumoniae* serotype 1 (2 x 10^6^ pfu) and euthanized 36 hpi. (**D**), Absolute numbers of neutrophils (CD45^+^ CD11b^+^ SiglecF^-^ F4/80^-^ Ly6G^+^) in BALF and lung parenchyma were evaluated by flow cytometry. Individual values and mean ± SD from three independent experiments are shown. Mann-Whitney test. (**E**), Representative dot plots (left panels) and absolute numbers (right panels) of TNF-α-producing-neutrophils (top panel) and -interstitial macrophages (bottom panel). Data are representative of three independent experiments. Mann-Whitney test. (**F**), B6-MAIT^CAST^ and B6-MAIT^CAST^ *Mr1*^-/-^ mice were i.n. infected with *S. pneumoniae* serotype 1 (5 x 10^5^ pfu). Levels of MPO in lung homogenates at 48 hpi. Individual values and means ± SD from four independent experiments are shown. Unpaired t test. ns, not significant; *, P < 0.05; **, P < 0.01 ; ****, P < 0.0001.

We next assessed whether MAIT cells could rather control antibacterial functions of phagocytes. We found more TNF-α-expressing neutrophils and macrophages in the lungs of pneumococcus-infected B6-MAIT^CAST^ mice than B6-MAIT^CAST^ *Mr1*^-/-^ mice (**Figure 5E**). Moreover, the levels of myeloperoxidase (MPO) were significantly reduced in the lung of B6-MAIT^CAST^ *Mr1*^-/-^ mice (**Figure 5F**). Notably, neutrophil frequency and MPO levels were also positively correlated in infected B6-MAIT^CAST^ mice but not in B6-MAIT^CAST^ *Mr1*^-/-^ mice (**Figure S5F**), suggesting dysregulated activity of neutrophils in absence of MAIT cells. Together, these findings demonstrate that MAIT cells restrain the growth and dissemination of *S.p.* likely through the control of myeloid cells’ activity.

### Prophylactic 5-OP-RU treatment protects mice from pneumococcus-induced lethal pneumonia

MAIT cells have emerged as appealing targets for cell-based immune intervention^8,12^ including during lung infections^20,42^. To evaluate whether MAIT cell stimulation using 5-OP-RU could provide protection against *S.p.*-mediated lethal pneumonia, we instilled MAIT cell cognate Ag intranasally. We observed that 5-OP-RU induced a decrease in MAIT cell frequency in a dose-dependent manner (**Figure 6A**). However, the absolute count of MAIT cells was only reduced at the highest dose (**Figure 6A**), an effect potentially due to TCR down-modulation in response to over-activation (**Figure S5A**). Lung MAIT cells also expressed CD69 and PD-1 in a dose-dependent manner (**Figure 6B**). Activated MAIT cells produced IL-17A in a dose-dependent manner (**Figure 6C**) but produced little to no IFN-γ (**Figure 6C**). Of note, unlike previous observations using combination of 5-OP-RU and TLR ligands, no change in Ki67 expression was observed, which suggests no early proliferation for MAIT cells upon standalone 5-OP-RU instillation (**Figure S5B**). Moreover, lung MAIT cells from 5-OP-RU-experienced mice were still able to respond (*e.g.* CD69 and IL-17A production) to a secondary challenge with 5-OP-RU (**Figure S5C**), suggesting absence of long-term hyporesponsiveness upon primary TCR triggering.

**Figure 6:**
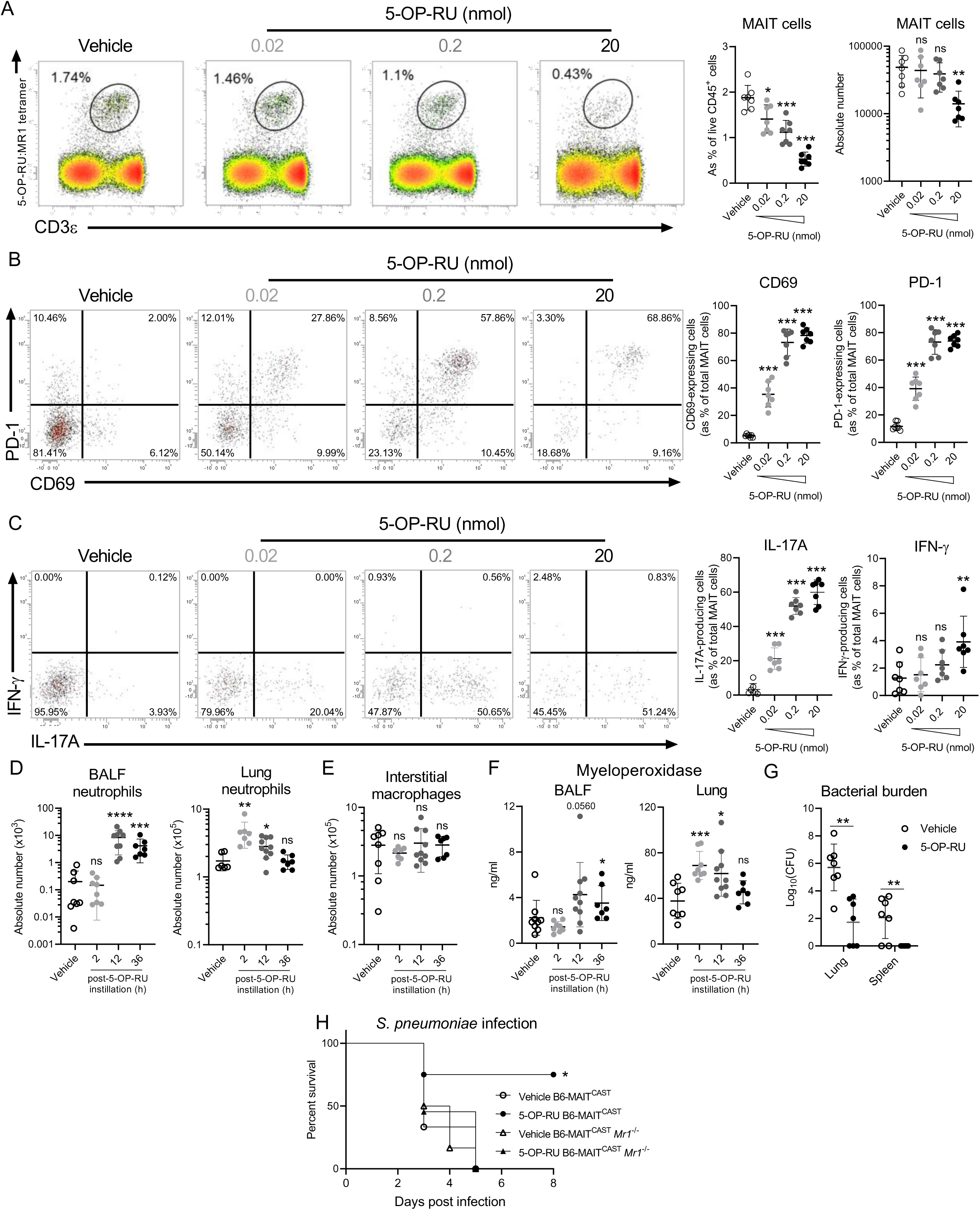
Effect of 5-OP-RU on host response to pneumococcus-induced lethal pneumoniae. B6-MAIT^CAST^ and B6-MAIT^CAST^ *Mr1*^-/-^ mice were i.n. treated once with indicated doses of 5-OP-RU or vehicle as control. (**A-C**) B6-MAIT^CAST^ mice were euthanized 10h post-instillation. (**A**), Representative dot plots of MAIT cell frequency in lungs of treated mice are shown in the left panel. Individual values and mean ± SD of MAIT cell frequency and absolute counts from two independent experiments are shown in the right panel. Mann-Whitney test. (**B**), Expression of activation markers by lung MAIT cells. Representative dot plots of CD69 and PD-1 expression on MAIT cells are shown in the left panel. Individual values and mean ± SD of CD69- and PD-1-expressing MAIT cells from two independent experiments are shown in the right panel. Mann-Whitney test. (**C**), Production of cytokines by lung MAIT cells. Representative dot plots of IL-17A and IFN-γ expressions by MAIT cells are shown in the left panel. Individual values and mean ± SD of IL-17A- and IFN-γ-expressing MAIT cells from two independent experiments are shown in the right panel. Mann-Whitney test. (**D-I**), B6-MAIT^CAST^ mice were treated i.n. with 0.2 nmol of 5-OP-RU and were euthanized at indicated time points. (**D, E**), Relative proportions and total numbers of neutrophils (**D**) and interstitial-like macrophages (**E**) were determined in BALF and/or lung. Individual values and mean ± SD from three independent experiments are shown. Mann-Whitney test. (**F**), Levels of MPO in BALF and lung homogenates. Individual values and mean ± SD of three independent experiments are shown. Mann-Whitney test. (**G-H**), Ten hours post-instillation, vehicle- or 5-OP-RU-treated mice were infected with 2 x 10^6^ pfu of pneumococcus serotype 1. (**G**), Bacterial burden was evaluated in total lung and spleen at 60 hpi. Individual values and mean ± SD from two experiments are shown. Mann-Whitney test. (**H**), Survival of mice was monitored daily. Data are pooled from four independent experiments. Mantel-Cox test. ns, not significant; *, P < 0.05; **, P < 0.01 ; ***, P < 0.001; ****, P < 0.0001.

Since 0.2 nmol of 5-OP-RU was sufficient to activate MAIT cells, we used this dose for the rest of the study. Consistent with the enrichment in MAIT17 cells in the lungs, 5-OP-RU-treated B6-MAIT^CAST^ mice displayed an increased number of neutrophils (**Figure 6D**) but not of interstitial macrophages (**Figure 6E**). In line with the lung neutrophilia, increased myeloperoxidase release (**Figure 6F**) was detected in 5-OP-RU-treated mice. Thus, we evaluated the outcome of 5-OP-RU prophylactic treatment on bacterial containment and survival during *S. pneumoniae*-induced lethal pneumonia. Intranasal 5-OP-RU administration led to reduced bacterial burdens in the lungs and limited systemic dissemination (**Figure 6G**). Moreover, 5-OP-RU administration enhanced survival in B6-MAIT^CAST^, but not in B6-MAIT^CAST^ *Mr1*^-/-^ mice (**Figure 6H**). Altogether, this demonstrates that TCR-mediated activation of lung MAIT cells induces a T_H_17-like innate response that protects against *Streptococcus pneumoniae*-induced lethal pneumonia.

## Discussion

MAIT cells are key actors in host response to Gram-negative bacteria, yet their contribution in defence against Gram-positive bacteria remains poorly defined. This question is of upmost importance given that *S. pneumoniae* is one the leading cause of CAP worldwide and culminating in 500,000 deaths per year^43^. Here, we found that MAIT cells confer protection to pneumococcus-related lung infection. Protective activity relies on the ability of MAIT cells to control bacterial outgrowth likely through the regulation of neutrophil and macrophage functions. Importantly, high content in MAIT cells (B6-MAIT^CAST^) confers an advantage in host resistance to pneumococcal infection as compared to classical B6 mice and MAIT cell-deficient mice that likely explains the absence of MAIT cell-dependent protection recently reported in a B6 setting^7^.

Biological assays and transcriptional analyses indicated that lung MAIT cells are stimulated by cytokines and TCR-mediated signals to rapidly acquire antibacterial functions upon pneumococcal infection. Our data show that cytokines initially drive MAIT cell activation during infection, and that TCR engagement might occur later on during the course of infection, which could be related to antigen bioavailability. Indeed, pneumococcus is initially rapidly controlled, followed by a sharp growth phase. Interestingly, our data suggest that several pro-inflammatory cytokines participate in MAIT cell activation, including IL-12 and/or IL-23, IL-15, IL-18 and type I IFN. This is consistent with previous reports in the context of respiratory infections with Influenza A virus^44^, *Salmonella typhimurium*^42^ and *Klebsiella pneumoniae*^20^. Our data also suggest a role for IL-1 similar to previous observation in the skin^45^. Whether the protective functions of lung MAIT cells are supported by cognate and/or cytokine signals remain to be determined.

This observation also raises the question about the involvement of redundant *vs* non-redundant pathways in the MAIT cell-dependent beneficial effect. For instance, MAIT cells promptly produce IL-17A but appear to be dispensable for neutrophil recruitment. This may be explained by a functional redundancy with lung γδT17 cells, which are critical for neutrophil recruitment^5–7^. Indeed, γδT17 cells outnumber MAIT17 cells in the lungs of B6-MAIT^CAST^ mice. Thus, this numerical superiority likely offers an advantage to γδT17 cells over MAIT cells to locally sense activating cytokines such as IL-1β or IL-23^5^.

The contribution of the MAIT-derived effector molecules participating in the protective effect remains to be determined. Transcriptomic analyses indicate the overexpression or induction of many genes in MAIT cells from infected mice previously shown to control neutrophil and/or macrophage functions including IFN-γ^46^, IL-22^47^, TNF-α^48,49^, lymphotoxin-α/β^50,51^, MIF^52^ and GM-CSF^53^. Understanding the respective role of these MAIT cell-dependent factors in myeloid cells’ effector functions will be of interest in the design of new immune-activators able to control *S. pneumoniae* infection.

Moreover, this study also reveals that prophylactic TCR-mediated MAIT cell activation is sufficient to protect mice against lethal *S. pneumoniae* infection. Indeed, local administration of 5-OP-RU - in absence of combinatorial mucosal adjuvants - results in a rapid activation of lung MAIT cells (2h post-instillation) culminating in the production of copious amounts of IL-17A. This is paralleled with a rapid recruitment of neutrophils, which is associated with a better containment of the bacteria in the lung tissue and complete abrogation of bacterial dissemination. While this likely explains the anti-pneumococcal activity, it will be of interest to decipher the overall molecular and cellular pathways of the innate immune cascade elicited by 5-OP-RU to develop optimized protocols. Since our lab has already highlighted a similar protective axis using the iNKT cell ligand, α-galactosylceramide - in combination or not with IL-7^4,6^, understanding the similarities and discrepancies of the overall response induced by TCR triggering of iNKT *vs* MAIT cells will be of upmost importance to optimize antibacterial strategies based on the harnessing of innate-like T cell functions.

Our clinical data revealed a decreased frequency of circulating MAIT cells in ICU patients with severe pneumococcal pneumonia; a phenomenon associated with a phenotype of highly activated cells. This decrease may suggest a migration of activated MAIT cells into the diseased tissue as already suggested during SARS-CoV-2 infections^32^ and ulcerative colitis^54,55^. Moreover, the level of activation appears to correlate with key clinical features, an observation reminiscent with our previous report in patients with severe COVID-19^32^. Since patients with highly activated MAIT cells display lower hypoxemia, it is tempting to speculate that MAIT cells also have a beneficial effect in severe pneumococcal infections. The experimental model points towards a key role for MAIT cells in regulating the functions of neutrophils and interstitial macrophages. Whether this MAIT cell-dependent contribution in disease resistance is relevant in ICU patients with pneumococcal CAP remains, however, uncertain given the rapid introduction of antibiotic treatment. Nonetheless, this antibacterial activity may play a significant role in pauci-symptomatic or mild patients. It is also noteworthy that lung MAIT cells from *S. pneumoniae*-infected mice acquire a tissue repair signature, an important immune function in the context of severe pneumonia and ARDS^56^.

In conclusion, our study unravelled that lung MAIT cells regulate myeloid cell’s activity as well as improve bacterial control and limit pneumonia. Moreover, exogenous TCR-triggering of MAIT cells is sufficient to protect mice against lethal pneumococcal pneumonia. Altogether, these findings should stimulate further studies on MAIT cells in severe pneumonia and may be valuable for the design of innovative therapeutic options.

## Methods

### Mice

All mice were housed and bred under specific pathogen free conditions in the animal facility of Tours University. Inbred C57BL/6JRj (B6) mice were purchased from Janvier (Le Genest-St-Isle, France). B6-MAIT^CAST^ ^35^, B6-MAIT^CAST^ *Mr1*^-/-^ and *Nr4a1*-GFP B6-MAIT^CAST^ ^39^ mice were kindly provided by Dr. O. Lantz (Institut Curie, Paris, France). Sex-matched 8-12 week-old mice were used in the study. All animal work was conformed to the French governmental and committee guidelines for the care and use of animals for scientific purposes; and was approved by the national ethic committee under APAFIS approval numbers #4885-201604071220401 and #37373-2022051211235184.

### Clinical study, patient population and approval

All biological samples were collected from adult patients admitted in ICU (University Hospital of Tours) with diagnosed bacterial pneumonia (n = 19). All patients or their next of kin gave their consent for enrolment in the study. This work is part of the clinical study ImPACT (NCT03379207) that was approved by the ethic committee “Comité de Protection des Personnes Ile-de-France 8” under the agreement number 2017-A01841-52. Blood samples from healthy donors (matched in sex and age) were collected from the “Etablissement Français du Sang” (CPDL-PLER-2019 188).

### Reagents, cell lines and Abs

Monoclonal Abs against mouse CD45 (30F11), CD3 (145-2C11), CD19 (1D3), CD69 (H1.2F3), TCRδ (GL3), CD11b (M1/70), Ly6G (1A8), SiglecF (E50-2440), F4/80 (BM8), IFN-γ (XMG1.2), IL-17A (TC11-18H10-1), Ki67 (11F6) and isotype controls were used. Monoclonal Abs against human CD45 (2D11), TCR Vα7.2 (3C10), CD161 (HP-3G10), IFN-γ (4S.B3), IL-17A (BL168) and isotype controls were used. All monoclonal antibodies were purchased from Biolegend (Amsterdam, The Netherlands), BD Pharmingen (Le Pont de Claix, France), Miltenyi-Biotec (Paris, France) or Thermo Fisher Scientific (Illkirch, France). Mouse and human MR1 tetramers were obtained from the National Institute of Allergy and Infectious Diseases Tetramer Facility (Emory University, Atlanta, GA). LIVE/DEAD® Stain kit was purchased from ThermoFisher Scientific. Anti-mouse (m)IL-1β (B122), anti-mIL-12p40 (C17.8), anti-mIL15 (AIO.3), anti-mIFNAR1 (MAR1-5A3), anti-mIL18 (YIGIF74-1G7) and isotype controls were from Bio X Cell (Lebanon, NH, USA).

### Pneumococcal strains, mutant generation and infection model

Clinical isolate of *S. pneumoniae* serotype 1 isolate E1586 sequence type ST304 has been described elsewhere^34^. The riboflavin-deleted mutants were derived from *S. pneumoniae* strain D39V^57^ and bacteria were grown and stored as previously described (**Table S3**). For transformation, D39V pneumococci were grown in C+ Y medium at 37°C, and subsequently treated with 100 ng/ml of synthetic CSP-1 for 10 min at 37°C to induce competence. The riboflavin operon was replaced by an erythromycin-resistance marker to construct the riboflavin operon-deficient mutant strain (Δ*ribDEHA::ery*). The upstream region and downstream region of riboflavin operon and the erythromycin marker were digested with the restriction enzyme SapI, then ligated. The ligation product was transformed into competent D39V bacteria. The complemented strain (Δ*ribDEHA::ribDEHA*) was constructed by integration of the pPEPZ plasmid, which integrates into the ZIP locus, with the entire *ribDEHA* + upstream riboswitch operon. Here, Golden Gate assembly using Esp3I was used with the upstream region and downstream region of the pPEPZ plasmid, riboflavin operon and a spectinomycin marker transformed into the D39VΔ*ribDEHA::ery* strain. Oligonucleotides used to amplify the various fragments are listed in **Table S3**. Transformants were selected by plating on Columbia agar blood plates supplemented with 2% (v/v) defibrinated sheep blood with either 0.5µg/ml erythromycin or 100µg/ml spectinomycin.

All mice were anesthetized and infected by the intranasal (i.n.) route with 50µl of PBS containing live *S. pneumoniae* strains. Survival and bacterial load experiments were performed using serotype 1 at 5.10^5^ CFU corresponding to a LD50 in B6 mice. All other experiments with serotype 1 were performed using a LD100 (2.10^6^ CFU). Mechanistic experiments using the D39V strains were performed at 4.10^6^ CFU. Assessment of 5-OP-RU protection was performed using serotype 1 at LD100. Clinical manifestations and survival were monitored daily following infection.

### Tissue collection and preparation of single cell suspensions

Lungs were collected at indicated time points and processed as previously described^6^. Briefly, lungs were perfused with 5 ml of PBS injected into the right ventricle, then harvested and processed using a gentleMACS dissociator (Miltenyi Biotec) in PBS containing 125µg/ml of Liberase (Roche; Meylan, France) and 100μg/ml of DNase type I (Roche). Homogenates were resuspended in PBS 2% FCS after passages through a 20G needle and filtered through a 100-µm cell strainer (Dutscher). Erythrocytes were lysed using RBC lysis buffer (Sigma-Aldrich) and suspensions were then passed through a 40-µm cell strainer. Pellets were resuspended in FACS buffer for further analyses. In some cases, bronchoalveolar lavage fluids (BALF) were also collected. BALF were centrifuged at 450g for 5 minutes and erythrocytes were removed using Red Blood Cell (RBC) lysis buffer. Cells were resuspended in FACS buffer containing PBS 2% FCS + 2 mM EDTA.

PBMC were isolated through density gradient centrifugation using Histopaque-1077 solution (Sigma-Aldrich). RBC were removed using RBC lysis buffer (Sigma-Aldrich).

### Assessment of bacterial burden

Mice were euthanized at indicated time point. Bacterial loads were evaluated in BALF, lung parenchyma and spleen. Serial dilutions of tissue homogenates (using gentleMacs dissociator) or BALF were next plated onto blood agar and incubated at 37°C in the presence of 5 % of CO_2_. Colony-forming units were enumerated 24 h later.

### In vitro activation of human MAIT cells

PBMC from healthy donors were cultured with medium or fixed pneumococcal strains at 50 MOI for 12 h. Glycerol stocks of pneumococcus were washed three times with PBS and bacteria were subsequently fixed in 4% of paraformaldehyde (30 min at 4°C). Three additional washing steps were performed prior culture. In some cases, 10 µg/ml of neutralizing antibodies were added to the culture.

### Flow cytometry

Surface markers were stained using monoclonal antibodies at the appropriate dilution at 4°C for 30 min. For intracellular staining, cell suspensions were pre-incubated at 37°C for 2h in complete RPMI medium supplemented with Monensin and Brefeldin A (Biolegend). Fixation buffer (Biolegend) was used, and cells were next permeabilized using intracellular staining permeabilization wash buffer (Biolegend) before intracellular staining. Cells were then analyzed on a MACSQuant Analyzer (Miltenyi-Biotec) and data analysis was performed using the VenturiOne software.

### 5-OP-RU preparation and instillation

5-OP-RU (5-A-RU/MeG) was prepared extemporaneously by mixing equal amounts of 5-A-RU (Cayman Chemical) and MeG (Sigma-Aldrich) in PBS and incubated for 15 min at RT prior intranasal instillation at the indicated amount in 50 µl of PBS.

### Single-cell RNA-seq and data pre-processing

Single-cell suspension of four lungs from naive or pneumococcus-infected (12 h and 36 hpi) mice were pooled for each condition, and MAIT cells (LiveDead^-^ CD45^+^ CD19^-^ CD3^+^ MR1:5- OP-RU tetramer^+^) were purified using a FACSMelody (BD Biosciences) (purity > 96%). Cells were counted under a microscope and 20,000 MAIT cells per condition were loaded onto a Chromium controller (10X Genomics). Reverse transcription and library preparation were performed using the Next GEM Cell 5’ Reagent Kits v2 according to the manufacturer’s protocol. Libraries were sequenced on a NovaSeq 6000 (Institut Curie, Paris). Cell Ranger Single-Cell Software Suite v7.1.0 was used for sample demultiplexing, barcode processing and single-cell gene counting using standards default parameters ^58^ and aligned to the reference genome *Mus musculus build mm10*.

### Single-cell RNA-seq analyses

#### Cell and gene filtering

Raw gene expression matrices generated per sample were passed to Seurat 5.1.0 for pre-processing. Low quality cells were filtered using ddqcR^59^. Cells expressing *Cd19* and/or *Cd68* were removed as non-MAIT cell contaminants. Then, basic workflow from Seurat 5.1.0^60^ was used. Moreover, transcripts expressed in less than three cells were removed. Cells considered as doublet were also removed using DoubletFinder^61^. Thus, we retained 13,116 cells for downstream analyses. Samples were next aggregated to compare cells across conditions. Uniform Manifold Approximation and Projection (UMAP) was applied using 19 principal components (k = 30; min.dist = 0.3), and eight clusters were identified using the Louvain algorithm with a resolution of 0.2.

#### Plots

All feature plots were created using scCustomize package 2.1.2. Differentially expressed genes were identified using FindAllMarkers. ClusterMap to evaluate similarities between clusters was created using scanpy^62^. The following signatures have been used: “Response of MAIT cells to acute lung infection”^40^ and “Tissue repair”^41^.

### Statistical analysis

All statistical analysis were performed using GraphPad Prism software. The statistical significance was evaluated using Student’s t test or ANOVA1 with a Bonferroni post-test, or their respective non-parametric tests Mann-Whitney U tests or Kruskal-Wallis as indicated in figure legends. Possibility to use parametric tests has been assessed by using pre-test to check whether the population is Gaussian and the variance is equal. Survival rates were analyzed using a log-rank (Mantel-Cox) test. Results with a P value of less than 0.05 were considered significant. ns: not significant; *, p < 0.05; **, p <0.01; ***, p < 0.001, ****, P < 0.0001.

## Abbreviations

hpi: hours post-infection
MAIT: Mucosal-associated invariant T
MHC: major histocompatibility complex
WT: wild-type;

## Acknowledgments

We thank the University of Tours animal facility (PST-A) for their excellent mouse husbandry. The Cytometry and Single-cell Immunobiology core facility (Inserm, UMS61 Analysis of Biological Systems) is also acknowledged for expertise and technical assistance. We acknowledge the generous support from the NIAID Tetramer Facility (Emory University, Atlanta, GA) for providing MR1 tetramers. Aurélie Darbois (Institut Curie) is acknowledged for technical assistance. We are grateful to all healthcare co-workers involved especially Aurélie Aubrey, Delphine Chartier and Véronique Siméon for their excellent management of patient samples and clinical data. Stephan Ehrmann, Pierre-François Dequin, Annick Legras, Denis Garot, Emmanuelle Mercier, Charlotte Salmon-Gandonnière, Laetitia Bodet-Contentin, Marlène Morisseau, Stephan Mankikian and Walid Darwiche are acknowledged for patient inclusions. We thank all the patients and their families for their trust and confidence in our work. This work was supported by the Institut National de la Santé et de la Recherche Médicale (Inserm), the University of Tours and the Agence Nationale de la Recherche (ANR-21-CE07-0053; ANR-22-CE14-0047; ANR-23-CE17-0064; ANR-24-CE15-2471). V.M. and J.W.V. were supported by Swiss National Science Foundation project grant 310030_200792. E.B. is a recipient of a doctoral fellowship from the Ministère de l’Enseignement supérieur et de la Recherche. C.S.G.G. is a recipient of a Procope Mobility Fellowship. P.K. is supported by the University of Gothenburg. A.H., Y.J., M.B., M.Si. and C.P. are supported by Inserm. L.G., G.I., R.L. and T.B. are supported by the University of Tours. O.L. and M.Sa. are funded by INSERM, Institut Curie, European Research Council (ERC-2019-AdG-885435) and ANR (ANR-20-CE15-0028-01, ANR-23-CE15-001901, ANR-22-CE17-002702 and ANR-10-IDEX-0001-02 PSL).

## Author contributions

E.B. performed experiments, analyzed data, and wrote the paper with the input of all authors. G.I. designed and performed the bioinformatic analysis and analyzed data. L.G., A.H., R.L., V.M., M.L., M.B., M.F., A.Ga., C.S.G.G. performed the experiments and analyzed data. Y.J., A.Gu., M.Si., P.K. and T.M. provided critical input. J.W.V., M.Sa. and O.L. provided key reagents and provided critical input. T.B. and C.P. **supervised the project, performed experiments, analyzed data, and wrote the paper with the input of all authors.**

## Declaration of interests

The authors declare no competing interests.

Further information and requests for resources and reagents should be directed to and will be fulfilled by the Lead Contact, Christophe Paget.

## References

1. GBD 2016 Lower Respiratory Infections Collaborators (2018). Estimates of the global, regional, and national morbidity, mortality, and aetiologies of lower respiratory infections in 195 countries, 1990-2016: a systematic analysis for the Global Burden of Disease Study 2016. Lancet Infect. Dis. 18, 1191–1210. 10.1016/S1473-3099(18)30310-4.

2. ARDS Definition Task Force, Ranieri, V.M., Rubenfeld, G.D., Thompson, B.T., Ferguson, N.D., Caldwell, E., Fan, E., Camporota, L., and Slutsky, A.S. (2012). Acute respiratory distress syndrome: the Berlin Definition. JAMA 307, 2526–2533. 10.1001/jama.2012.5669.

3. Koppe, U., Suttorp, N., and Opitz, B. (2012). Recognition of Streptococcus pneumoniae by the innate immune system. Cell. Microbiol. 14, 460–466. 10.1111/j.1462-5822.2011.01746.x.

4. Ivanov, S., Fontaine, J., Paget, C., Macho Fernandez, E., Van Maele, L., Renneson, J., Maillet, I., Wolf, N.M., Rial, A., Léger, H., et al. (2012). Key role for respiratory CD103(+) dendritic cells, IFN-γ, and IL-17 in protection against Streptococcus pneumoniae infection in response to α-galactosylceramide. J. Infect. Dis. 206, 723–734. 10.1093/infdis/jis413.

5. Hassane, M., Demon, D., Soulard, D., Fontaine, J., Keller, L.E., Patin, E.C., Porte, R., Prinz, I., Ryffel, B., Kadioglu, A., et al. (2017). Neutrophilic NLRP3 inflammasome-dependent IL-1β secretion regulates the γδT17 cell response in respiratory bacterial infections. Mucosal Immunol. 10, 1056–1068. 10.1038/mi.2016.113.

6. Hassane, M., Jouan, Y., Creusat, F., Soulard, D., Boisseau, C., Gonzalez, L., Patin, E.C., Heuzé-Vourc’h, N., Sirard, J.-C., Faveeuw, C., et al. (2020). Interleukin-7 protects against bacterial respiratory infection by promoting IL-17A-producing innate T-cell response. Mucosal Immunol. 13, 128–139. 10.1038/s41385-019-0212-y.

7. Murray, M.P., Crosby, C.M., Marcovecchio, P., Hartmann, N., Chandra, S., Zhao, M., Khurana, A., Zahner, S.P., Clausen, B.E., Coleman, F.T., et al. (2022). Stimulation of a subset of natural killer T cells by CD103+ DC is required for GM-CSF and protection from pneumococcal infection. Cell Rep. 38, 110209. 10.1016/j.celrep.2021.110209.

8. Salio, M. (2022). Unconventional MAIT cell responses to bacterial infections. Semin. Immunol. 61–64, 101663. 10.1016/j.smim.2022.101663.

9. Constantinides, M.G., and Belkaid, Y. (2021). Early-life imprinting of unconventional T cells and tissue homeostasis. Science 374, eabf0095. 10.1126/science.abf0095.

10. Klenerman, P., Hinks, T.S.C., and Ussher, J.E. (2021). Biological functions of MAIT cells in tissues. Mol. Immunol. 130, 154–158. 10.1016/j.molimm.2020.12.017.

11. Provine, N.M., and Klenerman, P. (2020). MAIT Cells in Health and Disease. Annu. Rev. Immunol. 38, 203–228. 10.1146/annurev-immunol-080719-015428.

12. Godfrey, D.I., Le Nours, J., Andrews, D.M., Uldrich, A.P., and Rossjohn, J. (2018). Unconventional T Cell Targets for Cancer Immunotherapy. Immunity 48, 453–473. 10.1016/j.immuni.2018.03.009.

13. Legoux, F., Salou, M., and Lantz, O. (2020). MAIT Cell Development and Functions: the Microbial Connection. Immunity 53, 710–723. 10.1016/j.immuni.2020.09.009.

14. Godfrey, D.I., Koay, H.-F., McCluskey, J., and Gherardin, N.A. (2019). The biology and functional importance of MAIT cells. Nat. Immunol. 20, 1110–1128. 10.1038/s41590-019-0444-8.

15. Lantz, O., and Legoux, F. (2019). MAIT cells: programmed in the thymus to mediate immunity within tissues. Curr. Opin. Immunol. 58, 75–82. 10.1016/j.coi.2019.04.016.

16. Corbett, A.J., Eckle, S.B.G., Birkinshaw, R.W., Liu, L., Patel, O., Mahony, J., Chen, Z., Reantragoon, R., Meehan, B., Cao, H., et al. (2014). T-cell activation by transitory neo-antigens derived from distinct microbial pathways. Nature 509, 361–365. 10.1038/nature13160.

17. Zhao, Z., Wang, H., Shi, M., Zhu, T., Pediongco, T., Lim, X.Y., Meehan, B.S., Nelson, A.G., Fairlie, D.P., Mak, J.Y.W., et al. (2021). Francisella tularensis induces Th1 like MAIT cells conferring protection against systemic and local infection. Nat. Commun. 12, 4355. 10.1038/s41467-021-24570-2.

18. Meierovics, A., Yankelevich, W.-J.C., and Cowley, S.C. (2013). MAIT cells are critical for optimal mucosal immune responses during in vivo pulmonary bacterial infection. Proc. Natl. Acad. Sci. 110, E3119–E3128. 10.1073/pnas.1302799110.

19. Wang, H., D’Souza, C., Lim, X.Y., Kostenko, L., Pediongco, T.J., Eckle, S.B.G., Meehan, B.S., Shi, M., Wang, N., Li, S., et al. (2018). MAIT cells protect against pulmonary Legionella longbeachae infection. Nat. Commun. 9, 3350. 10.1038/s41467-018-05202-8.

20. López-Rodríguez, J.C., Hancock, S.J., Li, K., Crotta, S., Barrington, C., Suárez-Bonnet, A., Priestnall, S.L., Aubé, J., Wack, A., Klenerman, P., et al. (2023). Type I interferons drive MAIT cell functions against bacterial pneumonia. J. Exp. Med. 220, e20230037. 10.1084/jem.20230037.

21. Kurioka, A., van Wilgenburg, B., Javan, R.R., Hoyle, R., van Tonder, A.J., Harrold, C.L., Leng, T., Howson, L.J., Shepherd, D., Cerundolo, V., et al. (2018). Diverse Streptococcus pneumoniae Strains Drive a Mucosal-Associated Invariant T-Cell Response Through Major Histocompatibility Complex class I-Related Molecule-Dependent and Cytokine-Driven Pathways. J. Infect. Dis. 217, 988–999. 10.1093/infdis/jix647.

22. Hartmann, N., McMurtrey, C., Sorensen, M.L., Huber, M.E., Kurapova, R., Coleman, F.T., Mizgerd, J.P., Hildebrand, W., Kronenberg, M., Lewinsohn, D.M., et al. (2018). Riboflavin Metabolism Variation among Clinical Isolates of Streptococcus pneumoniae Results in Differential Activation of Mucosal-associated Invariant T Cells. Am. J. Respir. Cell Mol. Biol. 58, 767–776. 10.1165/rcmb.2017-0290OC.

23. Riffelmacher, T., Paynich Murray, M., Wientjens, C., Chandra, S., Cedillo-Castelán, V., Chou, T.-F., McArdle, S., Dillingham, C., Devereaux, J., Nilsen, A., et al. (2023). Divergent metabolic programmes control two populations of MAIT cells that protect the lung. Nat. Cell Biol. 25, 877–891. 10.1038/s41556-023-01152-6.

24. Lu, B., Liu, M., Wang, J., Fan, H., Yang, D., Zhang, L., Gu, X., Nie, J., Chen, Z., Corbett, A.J., et al. (2020). IL-17 production by tissue-resident MAIT cells is locally induced in children with pneumonia. Mucosal Immunol. 13, 824–835. 10.1038/s41385-020-0273-y.

25. Hannaway, R.F., Wang, X., Schneider, M., Slow, S., Cowan, J., Brockway, B., Schofield, M.R., Morgan, X.C., Murdoch, D.R., and Ussher, J.E. (2020). Mucosal-associated invariant T cells and Vδ2+ γδ T cells in community acquired pneumonia: association of abundance in sputum with clinical severity and outcome. Clin. Exp. Immunol. 199, 201–215. 10.1111/cei.13377.

26. Grimaldi, D., Le Bourhis, L., Sauneuf, B., Dechartres, A., Rousseau, C., Ouaaz, F., Milder, M., Louis, D., Chiche, J.-D., Mira, J.-P., et al. (2014). Specific MAIT cell behaviour among innate-like T lymphocytes in critically ill patients with severe infections. Intensive Care Med. 40, 192–201. 10.1007/s00134-013-3163-x.

27. Ouyang, L., Wu, M., Shen, Z., Cheng, X., Wang, W., Jiang, L., Zhao, J., Gong, Y., Liang, Z., Weng, X., et al. (2021). Activation and Functional Alteration of Mucosal-Associated Invariant T Cells in Adult Patients With Community-Acquired Pneumonia. Front. Immunol. 12, 788406. 10.3389/fimmu.2021.788406.

28. Ouyang, L., Wu, M., Shen, Z., Cheng, X., Wang, W., Jiang, L., Zhao, J., Gong, Y., Liang, Z., Weng, X., et al. (2021). Activation and Functional Alteration of Mucosal-Associated Invariant T Cells in Adult Patients With Community-Acquired Pneumonia. Front. Immunol. 12. 10.3389/fimmu.2021.788406.

29. Jochems, S.P., Marcon, F., Carniel, B.F., Holloway, M., Mitsi, E., Smith, E., Gritzfeld, J.F., Solórzano, C., Reiné, J., Pojar, S., et al. (2018). Inflammation induced by influenza virus impairs human innate immune control of pneumococcus. Nat. Immunol. 19, 1299–1308. 10.1038/s41590-018-0231-y.

30. Aliberti, S., Dela Cruz, C.S., Amati, F., Sotgiu, G., and Restrepo, M.I. (2021). Community-acquired pneumonia. Lancet Lond. Engl. 398, 906–919. 10.1016/S0140-6736(21)00630-9.

31. Parrot, T., Gorin, J.-B., Ponzetta, A., Maleki, K.T., Kammann, T., Emgård, J., Perez-Potti, A., Sekine, T., Rivera-Ballesteros, O., Karolinska COVID-19 Study Group, et al. (2020). MAIT cell activation and dynamics associated with COVID-19 disease severity. Sci. Immunol. 5. 10.1126/sciimmunol.abe1670.

32. Jouan, Y., Guillon, A., Gonzalez, L., Perez, Y., Boisseau, C., Ehrmann, S., Ferreira, M., Daix, T., Jeannet, R., François, B., et al. (2020). Phenotypical and functional alteration of unconventional T cells in severe COVID-19 patients. J. Exp. Med. 217. 10.1084/jem.20200872.

33. Flament, H., Rouland, M., Beaudoin, L., Toubal, A., Bertrand, L., Lebourgeois, S., Rousseau, C., Soulard, P., Gouda, Z., Cagninacci, L., et al. (2021). Outcome of SARS-CoV-2 infection is linked to MAIT cell activation and cytotoxicity. Nat. Immunol. 10.1038/s41590-021-00870-z.

34. Muñoz, N., Van Maele, L., Marqués, J.M., Rial, A., Sirard, J.-C., and Chabalgoity, J.A. (2010). Mucosal administration of flagellin protects mice from Streptococcus pneumoniae lung infection. Infect. Immun. 78, 4226–4233. 10.1128/IAI.00224-10.

35. Cui, Y., Franciszkiewicz, K., Mburu, Y.K., Mondot, S., Le Bourhis, L., Premel, V., Martin, E., Kachaner, A., Duban, L., Ingersoll, M.A., et al. (2015). Mucosal-associated invariant T cell-rich congenic mouse strain allows functional evaluation. J. Clin. Invest. 125, 4171–4185. 10.1172/JCI82424.

36. Kiselev, V.Y., Kirschner, K., Schaub, M.T., Andrews, T., Yiu, A., Chandra, T., Natarajan, K.N., Reik, W., Barahona, M., Green, A.R., et al. (2017). SC3: consensus clustering of single-cell RNA-seq data. Nat. Methods 14, 483–486. 10.1038/nmeth.4236.

37. El Morr, Y., Fürstenheim, M., Mestdagh, M., Franciszkiewicz, K., Salou, M., Morvan, C., Dupré, T., Vorobev, A., Jneid, B., Premel, V., et al. (2024). MAIT cells monitor intestinal dysbiosis and contribute to host protection during colitis. Sci. Immunol. 9, eadi8954. 10.1126/sciimmunol.adi8954.

38. Legoux, F., Gilet, J., Procopio, E., Echasserieau, K., Bernardeau, K., and Lantz, O. (2019). Molecular mechanisms of lineage decisions in metabolite-specific T cells. Nat. Immunol. 20, 1244–1255. 10.1038/s41590-019-0465-3.

39. du Halgouet, A., Darbois, A., Alkobtawi, M., Mestdagh, M., Alphonse, A., Premel, V., Yvorra, T., Colombeau, L., Rodriguez, R., Zaiss, D., et al. (2023). Role of MR1-driven signals and amphiregulin on the recruitment and repair function of MAIT cells during skin wound healing. Immunity 56, 78–92.e6. 10.1016/j.immuni.2022.12.004.

40. Hinks, T.S.C., Marchi, E., Jabeen, M., Olshansky, M., Kurioka, A., Pediongco, T.J., Meehan, B.S., Kostenko, L., Turner, S.J., Corbett, A.J., et al. (2019). Activation and In Vivo Evolution of the MAIT Cell Transcriptome in Mice and Humans Reveals Tissue Repair Functionality. Cell Rep. 28, 3249–3262.e5. 10.1016/j.celrep.2019.07.039.

41. Linehan, J.L., Harrison, O.J., Han, S.-J., Byrd, A.L., Vujkovic-Cvijin, I., Villarino, A.V., Sen, S.K., Shaik, J., Smelkinson, M., Tamoutounour, S., et al. (2018). Non-classical Immunity Controls Microbiota Impact on Skin Immunity and Tissue Repair. Cell 172, 784–796.e18. 10.1016/j.cell.2017.12.033.

42. Wang, H., Kjer-Nielsen, L., Shi, M., D’Souza, C., Pediongco, T.J., Cao, H., Kostenko, L., Lim, X.Y., Eckle, S.B.G., Meehan, B.S., et al. (2019). IL-23 costimulates antigen-specific MAIT cell activation and enables vaccination against bacterial infection. Sci. Immunol. 4, eaaw0402. 10.1126/sciimmunol.aaw0402.

43. Bender, R.G., Sirota, S.B., Swetschinski, L.R., Dominguez, R.-M.V., Novotney, A., Wool, E.E., Ikuta, K.S., Vongpradith, A., Rogowski, E.L.B., Doxey, M., et al. (2024). Global, regional, and national incidence and mortality burden of non-COVID-19 lower respiratory infections and aetiologies, 1990–2021: a systematic analysis from the Global Burden of Disease Study 2021. Lancet Infect. Dis. 24, 974–1002. 10.1016/S1473-3099(24)00176-2.

44. van Wilgenburg, B., Loh, L., Chen, Z., Pediongco, T.J., Wang, H., Shi, M., Zhao, Z., Koutsakos, M., Nüssing, S., Sant, S., et al. (2018). MAIT cells contribute to protection against lethal influenza infection in vivo. Nat. Commun. 9, 4706. 10.1038/s41467-018-07207-9.

45. Constantinides, M.G., Link, V.M., Tamoutounour, S., Wong, A.C., Perez-Chaparro, P.J., Han, S.-J., Chen, Y.E., Li, K., Farhat, S., Weckel, A., et al. (2019). MAIT cells are imprinted by the microbiota in early life and promote tissue repair. Science 366. 10.1126/science.aax6624.

46. Esmann, L., Idel, C., Sarkar, A., Hellberg, L., Behnen, M., Möller, S., van Zandbergen, G., Klinger, M., Köhl, J., Bussmeyer, U., et al. (2010). Phagocytosis of apoptotic cells by neutrophil granulocytes: diminished proinflammatory neutrophil functions in the presence of apoptotic cells. J. Immunol. Baltim. Md 1950 *184*, 391–400. 10.4049/jimmunol.0900564.

47. Pavlidis, P., Tsakmaki, A., Pantazi, E., Li, K., Cozzetto, D., Digby-Bell, J., Yang, F., Lo, J.W., Alberts, E., Sa, A.C.C., et al. (2022). Interleukin-22 regulates neutrophil recruitment in ulcerative colitis and is associated with resistance to ustekinumab therapy. Nat. Commun. 13, 5820. 10.1038/s41467-022-33331-8.

48. Youn, C., Pontaza, C., Wang, Y., Dikeman, D.A., Joyce, D.P., Alphonse, M.P., Wu, M.-J., Nolan, S.J., Anany, M.A., Ahmadi, M., et al. (2023). Neutrophil-intrinsic TNF receptor signaling orchestrates host defense against Staphylococcus aureus. Sci. Adv. 9, eadf8748. 10.1126/sciadv.adf8748.

49. Kusnadi, A., Park, S.H., Yuan, R., Pannellini, T., Giannopoulou, E., Oliver, D., Lu, T., Park-Min, K.-H., and Ivashkiv, L.B. (2019). The Cytokine TNF Promotes Transcription Factor SREBP Activity and Binding to Inflammatory Genes to Activate Macrophages and Limit Tissue Repair. Immunity 51, 241–257.e9. 10.1016/j.immuni.2019.06.005.

50. Riffelmacher, T., Giles, D.A., Zahner, S., Dicker, M., Andreyev, A.Y., McArdle, S., Perez-Jeldres, T., van der Gracht, E., Murray, M.P., Hartmann, N., et al. (2021). Metabolic activation and colitis pathogenesis is prevented by lymphotoxin β receptor expression in neutrophils. Mucosal Immunol. 14, 679–690. 10.1038/s41385-021-00378-7.

51. Tan, T.Q. (2012). Pediatric invasive pneumococcal disease in the United States in the era of pneumococcal conjugate vaccines. Clin. Microbiol. Rev. 25, 409–419. 10.1128/CMR.00018-12.

52. Calandra, T., and Roger, T. (2003). Macrophage migration inhibitory factor: a regulator of innate immunity. Nat. Rev. Immunol. 3, 791–800. 10.1038/nri1200.

53. Hamilton, J.A. (2008). Colony-stimulating factors in inflammation and autoimmunity. Nat. Rev. Immunol. 8, 533–544. 10.1038/nri2356.

54. Haga, K., Chiba, A., Shibuya, T., Osada, T., Ishikawa, D., Kodani, T., Nomura, O., Watanabe, S., and Miyake, S. (2016). MAIT cells are activated and accumulated in the inflamed mucosa of ulcerative colitis. J. Gastroenterol. Hepatol. 31, 965–972. 10.1111/jgh.13242.

55. Serriari, N.-E., Eoche, M., Lamotte, L., Lion, J., Fumery, M., Marcelo, P., Chatelain, D., Barre, A., Nguyen-Khac, E., Lantz, O., et al. (2014). Innate mucosal-associated invariant T (MAIT) cells are activated in inflammatory bowel diseases. Clin. Exp. Immunol. 176, 266–274. 10.1111/cei.12277.

56. Thompson, B.T., Chambers, R.C., and Liu, K.D. (2017). Acute Respiratory Distress Syndrome. N. Engl. J. Med. 377, 562–572. 10.1056/NEJMra1608077.

57. Slager, J., Aprianto, R., and Veening, J.-W. (2018). Deep genome annotation of the opportunistic human pathogen Streptococcus pneumoniae D39. Nucleic Acids Res. 46, 9971–9989. 10.1093/nar/gky725.

58. Zheng, G.X.Y., Terry, J.M., Belgrader, P., Ryvkin, P., Bent, Z.W., Wilson, R., Ziraldo, S.B., Wheeler, T.D., McDermott, G.P., Zhu, J., et al. (2017). Massively parallel digital transcriptional profiling of single cells. Nat. Commun. 8, 14049. 10.1038/ncomms14049.

59. Subramanian, A., Alperovich, M., Yang, Y., and Li, B. (2022). Biology-inspired data-driven quality control for scientific discovery in single-cell transcriptomics. Genome Biol. 23, 267. 10.1186/s13059-022-02820-w.

60. Hao, Y., Stuart, T., Kowalski, M.H., Choudhary, S., Hoffman, P., Hartman, A., Srivastava, A., Molla, G., Madad, S., Fernandez-Granda, C., et al. (2024). Dictionary learning for integrative, multimodal and scalable single-cell analysis. Nat. Biotechnol. 42, 293–304. 10.1038/s41587-023-01767-y.

61. McGinnis, C.S., Murrow, L.M., and Gartner, Z.J. (2019). DoubletFinder: Doublet Detection in Single-Cell RNA Sequencing Data Using Artificial Nearest Neighbors. Cell Syst. 8, 329–337.e4. 10.1016/j.cels.2019.03.003.

62. Wolf, F.A., Angerer, P., and Theis, F.J. (2018). SCANPY: large-scale single-cell gene expression data analysis. Genome Biol. 19, 15. 10.1186/s13059-017-1382-0.

