## Supplementary Figures for "MAIT cells protect in severe pneumococcal pneumonia by regulating neutrophil/macrophage antimicrobial activities"

Barsac et al.,

**Supplementary Figures**

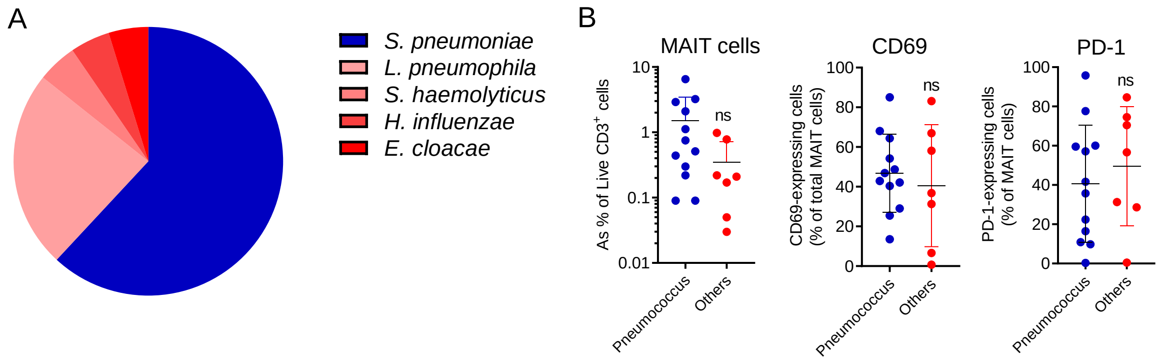


**Figure S1: Aetiology of bacterial CAP and activation status of circulating MAIT cells in patients according to the pathogen involved.** (**A**), Relative proportion of pathogens responsible for bacterial CAP in patients included in the study. (**B**), Flow cytometry analyses of MAIT cells in the blood of patients with pneumococcal- (n =12, blue) and non-pneumococcal (n = 7, red) bacterial CAP. Individuals and means ± SD of MAIT cell frequency, relative proportion of CD69 and PD-1-expressing MAIT cells are shown. Mann-Whitney test. ns, not significant.

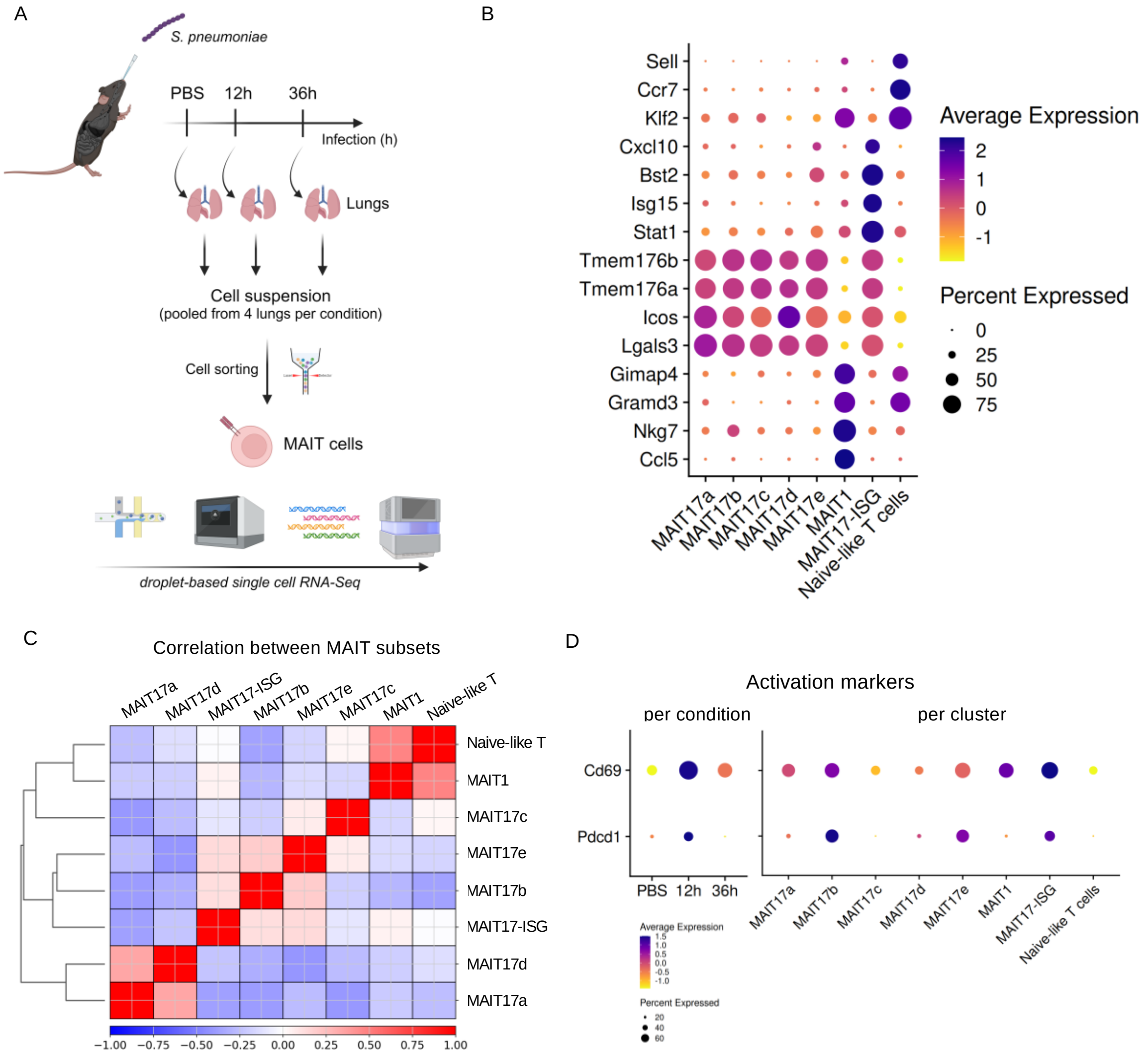


**Figure S2: Transcriptional signature of lung MAIT cells during pneumococcal infection.** (**A**), Experimental workflow of the single-cell RNA sequencing experiment. (**B**), Dot map showing expression of published marker genes of MAIT subsets[38] in each cluster. Circle size represents percentage of cells expressing each gene and color gradient indicates relative expression values. (**C**), Clustermap of MAIT clusters comparing each pair of clusters using the Pearson's correlation with hierarchical clustering. (**D**), Dot maps showing *Cd69* and *Pdcd1* expressions per condition (left panel) and per cluster (right panel).

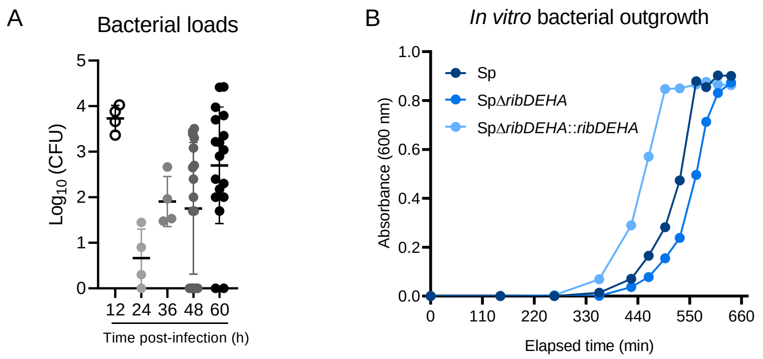


**Figure S3: Kinetics of pneumococcal strains' outgrowth.** (**A**), Kinetic evaluation of *in vivo* bacterial loads in BALF of B6-MAIT$^{CAST}$ mice upon *S. pneumoniae* infection (5 x $10^5$ pfu). Individual values and means ± SD from three independent experiments are shown. (**B**), *In vitro* growth curve of *S. pneumoniae* D39 parental and mutant strains in Todd Hewitt Broth medium containing 2% of yeast extracts.

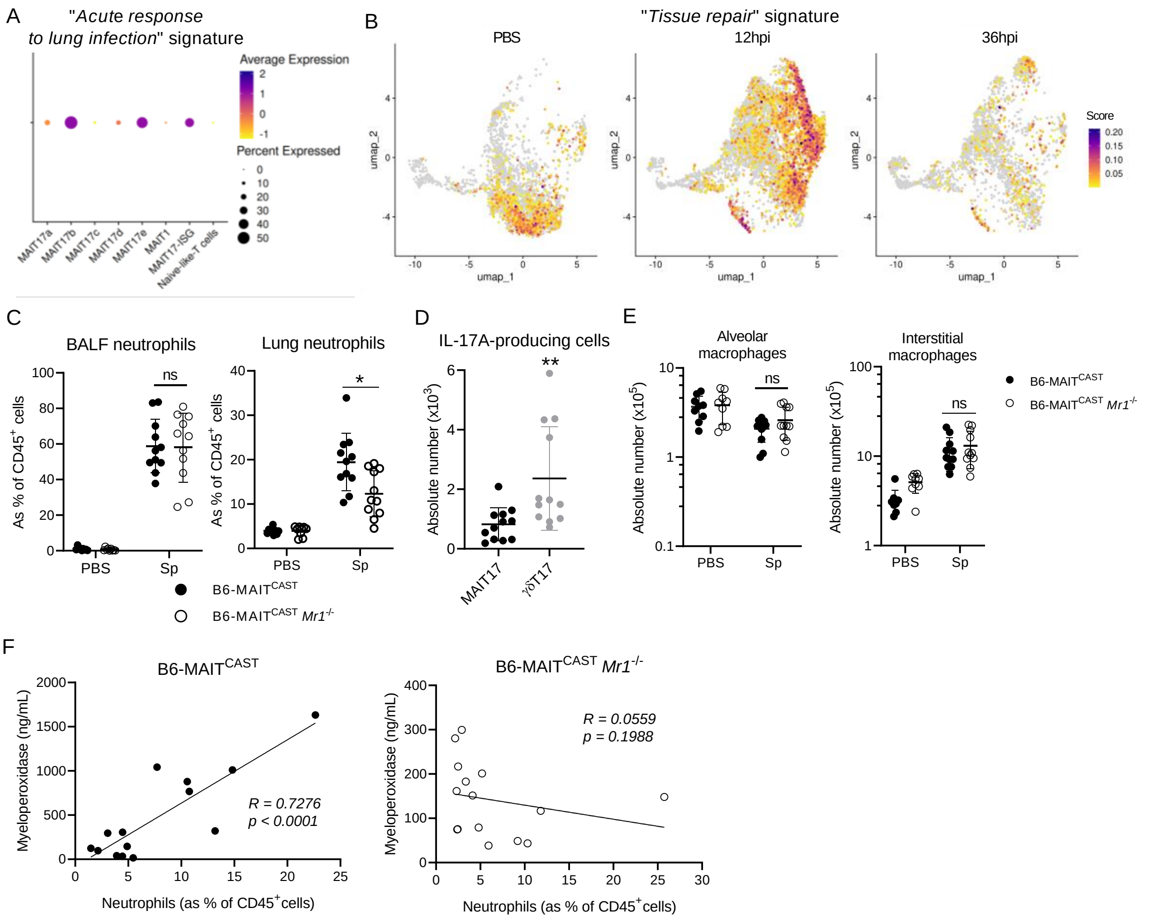


**Figure S4: Comparative analysis of the lung immune landscape in B6-MAIT$^{CAST}$ and B6-MAIT$^{CAST}$ *Mr1$^{-/-}$* mice infected with *S. pneumoniae*.** (**A**), Evaluation of the "*Acute response to lung infection*" signature on lung MAIT clusters. (**B**), Evaluation of "*Tissue repair*" signature on lung MAIT transcriptomes from naive or *S.p.*-infected mice. (**C-E**), B6-MAIT$^{CAST}$ and B6-MAIT$^{CAST}$ *Mr1$^{-/-}$* mice were i.n. infected with 2 x $10^6$ pfu of *S. pneumoniae serotype 1*. (**C**), Frequency of neutrophils (CD45$^+$ CD11b$^+$ SiglecF$^-$ F4/80$^-$ Ly6G$^+$) in BALF and lung parenchyma were evaluated by flow cytometry. Individual values and mean ± SD from three independent experiments are shown. Mann-Whitney test. (**D**), Absolute numbers of IL-17A-expressing MAIT cells and IL-17A-expressing γδT cells from the lungs of *S.p.*-infected B6-MAIT$^{CAST}$ mice. Individual values and mean ± SD of IL-17A-expressing cells from two independent experiments are shown. Mann-Whitney test. (**E**), Absolute numbers of alveolar macrophages (CD45$^+$ CD11$^{-/dim}$ SiglecF$^+$) and interstitial-like macrophages (CD45$^+$ CD11b$^+$ SiglecF$^-$ F4/80$^+$ Ly6G$^-$) were evaluated by flow cytometry. Individual values and mean ± SD from three independent experiments are shown. Mann-Whitney test. (**F**), Spearman's rank correlation of MPO levels and neutrophil frequencies in B6-MAIT$^{CAST}$ and B6-MAIT$^{CAST}$ *Mr1$^{-/-}$* mice infected with 5 x $10^5$ pfu of *S. pneumoniae* serotype 1. ns, not significant; *, $P < 0.05$; **, $P < 0.01$.

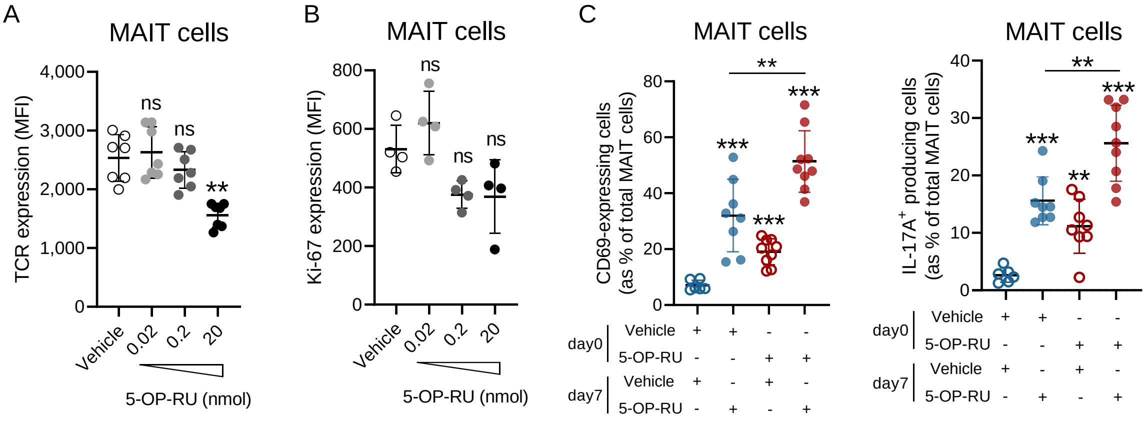


**Figure S5: TCR expression, proliferation and recall response of lung MAIT cells upon 5-OP-RU instillation.** B6-MAIT$^{CAST}$ mice were i.n. treated with indicated dose of 5-OP-RU or vehicle. (**A**), Expression of TCR on MAIT cells upon 5-OP-RU instillation (10h post-instillation) based on MR1 tetramer staining. Individual values and mean ± SD from two independent experiments are shown. Kruskal-Wallis test. (**B**), Expression of Ki67 in MAIT cells upon 5-OP-RU instillation (24h post-instillation). Individual values and mean ± SD are shown. Kruskal-Wallis test. (**C**), Activation status of lung MAIT cells upon 5-OP-RU instillation in naive or 5-OP-RU-experienced mice. Individual values and mean ± SD of CD69- (left panel) and IL-17A-expressing (right panel) MAIT cells from two independent experiments are shown. Mann-Whitney test. ns, not significant; **, $P < 0.01$ ; ***, $P < 0.001$.
