## Supplementary material for "MAIT cells protect in severe pneumococcal pneumonia by regulating neutrophil/macrophage antimicrobial activities": Table 1

| Characteristics | Overall  (n=19) | Pneumococcal CAP (n=12) | Non-pneumococcal CAP (n=7) |
| --- | --- | --- | --- |
| **Age (year)**, median (IQR) | 63 (58; 69) | 64 (58.8; 71.2) | 60 (48; 64) |
| **Male/Female**, n/n (%) | 10/9 (52.6%) | 6/6 (50%) | 4/3 (57.1%) |
| **BMI (kg/m2)**, median (IQR) | 28 (22; 30) | 25.5 (19.8; 29.7) | 28 (22; 30) |
| **Type 2 diabetes**, n (%) | 6 (31.6%) | 4 (33.3%) | 2 (28.6%) |
| **Hypertension**, n (%) | 8 (42.1%) | 6 (50%) | 2 (28.6%) |
| **Chronic respiratory disease**, n (%) | 7 (36.8%) | 5 (41.7%) | 2 (28.6%) |
| **Chronic kidney disease**, n (%) | 1 (5.3%) | 1 (8.3%) | 0 |
| **Chronic cardiovascular disease**, n (%) | 3 (15.8%) | 3 (25%) | 0 |
| **SAPS2**, median (IQR) | 45 (24; 58) | 40.5 (22.5; 56.3) | 46 (45; 74) |
| **SOFA at inclusion**, median (IQR) | 4 (2; 10) | 2.5 (2; 4.8) | 10 (4; 12) |
| **Invasive mechanical ventilation at inclusion** n (%) | 9 (47.4%) | 5 (41.7%) | 4 (57.1%) |

**Table 1 : Patients’ characteristics.** Quantitative data are reported as the median value and interquartile range; qualitative value are reported as n (%). BMI : body mass index; SAPS2 **:** Simplified Acute Physiology Score 2 ; SOFA : Sequential Organ Failure Assessment
